# Structural and Evolutionary Analysis of Human Growth Hormone Variants

**DOI:** 10.1101/2025.03.19.644177

**Authors:** Sonia Verma, Amit V Pandey

**Author notes:** (S.V.). (AVP).

## Abstract

Human growth hormone (GH) exerts its pleiotropic effects by binding to its receptor (GHR), leading to receptor dimerization and activation. We combined structural, evolutionary, and genetic analyses to elucidate the critical determinants of GH-GHR interaction and the impact of disease-causing mutations. Protein contact analysis revealed the specific amino acid residues involved in two distinct binding interfaces between GH and two chains of GHR. ConSurf analysis demonstrated significant sequence conservation in the receptor-binding regions of GH across species, highlighting their functional importance. A comprehensive list of known disease-causing mutations in GH was compiled and mapped to these binding interfaces and conserved regions. Computational site-directed mutagenesis (SDM) analysis predicted the impact of several mutations on protein stability, revealing both stabilizing and destabilizing effects. Sequence comparisons with orthologs from various species further supported the evolutionary conservation of key functional residues. Integrated analysis of contact residues between GH and GHR showed a strong correlation between receptor-binding residues, evolutionary conservation, and the occurrence of disease-associated mutations. These findings underscore the critical role of specific GH residues in mediating high-affinity interaction with its receptor, and how mutations in these conserved contact points can disrupt binding affinity and/or protein stability, ultimately leading to growth disorders. This multi-faceted approach provides valuable insights into the molecular mechanisms underlying growth hormone deficiency and related syndromes.

## 1. Introduction

Human growth and development is a highly regulated process governed by a complex network of steroid and peptide hormones. Steroid hormones, encompassing androgens (e.g., testosterone) and estrogens (e.g., estradiol), fulfill a critical role in growth, notably during puberty. These hormones contribute to the pubertal growth acceleration, facilitate bone maturation and epiphyseal closure (ultimately terminating linear growth), and are responsible for the development of secondary sexual characteristics and defect in genes encoding enzymes linked to steroid production lead to growth and developmental disorders [1–5]. Human growth hormone (GH), a peptide hormone, is critical in the complex regulation of growth, development, and metabolic processes throughout the human lifespan [6]. Growth hormone (GH), is principally synthesized by the anterior pituitary gland and serves as a pivotal regulator of linear growth, particularly during childhood and adolescence. GH manifests direct effects on diverse tissues, including the stimulation of osseous and cartilaginous growth, and indirect effects via the promotion of insulin-like growth factor-1 (IGF-1) production, primarily in the liver, which further stimulates growth. Steroid hormones can modulate the GH-IGF-1 axis by influencing GH secretion and IGF-1 production, underscoring the intricate hormonal regulation of human growth. The release of GH from the pituitary is induced by growth hormone-releasing hormone (GHRH), which originates in the hypothalamus and interacts with the GHRH receptor (GHRHR) on pituitary cells. GH subsequently interacts with the GH receptor (GHR) expressed on the surface of target cells throughout the body to elicit its multifaceted effects.

The gene encoding human growth hormone (GH1) is located on the long arm of chromosome 17, specifically at the 17q22-24 region [7]. This gene resides within a cluster of five closely related genes, including chorionic somatomammotropin hormone 1 (CSH1), CSH2, CSH-like 1 (CSHL1), and growth hormone variant 2 (GH2). This gene cluster spans approximately 65 kilobases [8]. The GH2 gene encodes a growth hormone variant expressed in the placenta, suggesting a specialized role during pregnancy. The clustering of these related genes implies potential for coordinated regulation and shared evolutionary origins. The mature hGH protein, comprising 191 amino acids, is derived from a larger 217-amino acid precursor through the cleavage of a 26-amino acid signal peptide [9–11]. The three-dimensional structure of hGH is characterized by a distinctive four-helix bundle motif, a structural arrangement crucial for its interaction with the GHR, stabilized by two intramolecular disulfide bonds (Cys53–Cys165, Cys182–Cys189) [12]. Comparative analysis reveals significant conservation of the hGH amino acid sequence across species, particularly within regions involved in GHR binding.

Synthesis and secretion of GH are regulated by somatotrophic cells located in the anterior pituitary gland [8, 9]. The pituitary gland, a small yet vital endocrine gland situated at the base of the brain, is under the direct influence of the hypothalamus which plays a central role through GHRH, which stimulates GH release, and somatostatin, which inhibits it. Ghrelin, produced in the gastrointestinal tract, also promotes GH production and release [13, 14]. The secretion of GH follows a pulsatile pattern, with the most significant release occurring during slow-wave sleep [15, 16]. Various factors influence this release, including circadian rhythm, sleep-wake cycles, stress levels, physical exercise, and nutritional status [17–21]. Following secretion, GH circulates in the bloodstream primarily bound to a growth hormone-binding protein (GHBP), which is essentially the cleaved extracellular domain of the GHR [22, 23]. GHBP acts as an GH reservoir and may modulate GH signaling. Its production is upregulated by GH, suggesting a negative feedback mechanism [24].

Beyond its established role in promoting growth, GH exerts diverse pleiotropic effects. Functions of GH extend beyond childhood and adolescence, maintaining tissue and organ health throughout adulthood. Specifically, GH influences somatic growth, contributes to bone elongation, and promotes muscle mass development. GH is also important for the regulation of essential metabolic pathways involving carbohydrates and lipids, facilitating protein synthesis, modulating the immune system. and maintaining proper bone density. The secretion of GH is pulsatile, with a significant surge during deep sleep phases, particularly slow-wave sleep [25]. Various internal and external factors, including sex, age, body fat composition, sleep patterns, stress levels, dietary intake, and physical activity, can significantly influence GH secretion. This dynamic responsiveness underscores the intricate regulation of GH in maintaining physiological equilibrium.

The diverse actions of GH are initiated through its binding to the growth hormone receptor (GHR), a transmembrane protein located on the surface of various target cells. These receptors are widely distributed throughout the body, including the liver, bone, and muscle tissues, as well as fat, kidney, brain, and skin. The GHR belongs to the cytokine receptor superfamily. The interaction between hGH and its receptor triggers a cascade of intracellular signaling events, notably the activation of the Janus kinase-signal transducer and activator of transcription via the JAK2/STAT5 signaling pathway. Upon GH binding, JAK2, a tyrosine kinase associated with the receptor, undergoes auto-phosphorylation and subsequently phosphorylates tyrosine residues on the GHR [26, 27]. These phosphorylated residues serve as docking sites for STAT family of transcription factors, primarily STAT-5. Once phosphorylated by JAK2, STAT proteins dissociate from the receptor, translocate to the nucleus, and activate the transcription of genes responsive to hGH [28]. A crucial downstream effect of hGH signaling is the production of IGF-1. The liver is the primary site of IGF-1 secretion in response to GH binding to its receptors on hepatocytes. IGF-1 acts on type 1 IGF receptors in various tissues to stimulate linear growth. Notably, IGF-1 also inhibits further GH release from the pituitary gland, establishing a negative feedback loop. While many of GH’s growth-promoting effects are mediated through IGF-1, GH also exerts direct effects on various physiological processes. It directly stimulates bone growth by acting on chondrocytes and osteoblasts, promotes protein synthesis by enhancing nitrogen retention, and influences glucose homeostasis, generally counteracting insulin’s effects. Additionally, GH stimulates the immune system and promotes fat breakdown by stimulating triglyceride breakdown and oxidation in adipocytes. These direct actions underscore GH’s multifaceted role in regulating growth and metabolism.

Disruptions in the intricate GH signaling pathway, often due to mutations in the GH1 gene, can lead to various growth disorders [29–31]. Isolated growth hormone deficiency (IGHD), characterized by insufficient GH production, results in short stature and metabolic abnormalities [32–36]. IGHD can arise from mutations in either the GH1 gene or the growth hormone-releasing hormone and its receptor (GHRH and GHRHR) genes [37]. Several types of IGHD exist, each with distinct genetic underpinnings and varying severity [6, 38]. Laron syndrome represents a primary form of GH insensitivity caused by GHR gene mutations. Mutations within the GH1 gene can lead to reduced hGH secretion or the production of variant GH proteins with impaired receptor binding or compromised signaling capabilities.

This study aims to provide an analysis of the human growth hormone protein, focusing on the interplay between its structure, evolutionary conservation, and the impact of disease-causing mutations on its interaction with the GHR. By integrating structural analyses, evolutionary conservation patterns, and the predicted effects of mutations, we seek to identify key determinants of GH function and understand the molecular mechanisms underlying GH-related disorders.

## 2. Materials and Methods

### 2.1 Sequence Alignments

The GH protein sequences were retrieved from the UniProt protein database and aligned using CLC Protein Workbench (www.qiagenbioinformatics.com). The Multiple Sequence alignment included sequences of Growth hormones from *Homo sapiens* (P01241), *Macaca mulatta* (P33093), *Rattus norvegicus* (P01244), *Mus musculus* (P06880), *Equus caballus* (P01245), *Sus scrofa* (P01248), *Bos Taurus* (P01246), *Ovis aries* (P67930), *Cavia porcellus* (Q9JKM4), *Meleagris gallopavo* (P22077), *Gallus gallus* (P08998), *Struthio camelus* (Q9PWG3), *Anguilla japonica* (P08899), *Carassius auratus* (O93359), and *Salmo salar* (Q5SDS1).

### 2.2 Evolutionary conservation of amino acids

The HGH protein sequence (NP_000506.2) was used as a query sequence to search the homologous sequences in the Uniref90 database by performing a PSI-Blast search [39]. The selected HGH sequences were aligned using CLUSTALW. The multiple sequence alignment (MSA) was used as input in the Consurf analysis, which calculates evolutionary (functional and structural) conserved residues in the protein based on the phylogenetic and structural similarity between sequences from different species [40]. The conservation score for each residue position in MSA represents the conservation of the residue.

### 2.3 Protein structure loop modeling

HGH crystal structure solved with its receptor (PDB Id: 3HHR, dimeric receptor) [12] has few missing residues in the loop regions (Chain A: 148-154; Chain B & C: 57-62, 73-78), those were modeled by using loop modeling module of YASARA [41]. For each loop region, ten loop conformations were generated and out of those, an energy minimized conformation was selected for the loop replacement. The structure models were visualized by Pymol (www.pymol.org) and rendered as ray-traced images with POVRAY (www.povray.org)

### 2.4 HGH-receptor interaction analysis

HGH structure (PDB Id: 3HHR, Chain A: HGH, Chains B & C: Receptor) was analyzed to find the residues, in contact with its receptor via covalent or non-covalent bonding. HGH-receptor contact map was created by submitting our modeled structure into PDBsum Generate (www.ebi.ac.uk/pdbsum) [12].

### 2.5 Disease-causing mutations

The data related to disease-causing mutations were obtained from UniProtKB [42]. UniProtKB compiles disease-causing mutation data related to a protein under-involvement in the disease section.

### 2.6 Prediction of protein stability upon mutation using site-directed mutagenesis tool (SDM)

SDM software was used for predicting the effect of selected human GH mutants on the stability of the protein and their potential as a disease-causing mutant [43] [44]. SDM software uses environment-specific substitution frequencies within homologous protein families to calculate a stability score. SDM software calculates a pseudo delta G score (stability score) for the selected mutants upon providing HGH structure and a list of selected mutant variants [45] [46]

### 2.7 Analysis of HGH allele frequencies from genome sequencing

Frequencies from Exome Aggregation Consortium [47], NHLBI Exome Sequencing Project (NHLBI Exome Sequencing Project, 2016), and 1000 Genomes [48] were collected and assigned to the main categories of the 1000 Genomes project.

## 3. Results

### 3.1 Analysis of GH sequence across species

The sequence comparison reveals varying degrees of identity and similarity between human growth hormone and its orthologs in different species (Table 1). As expected, the highest sequence identity (96%) and similarity (97%) are observed with Rhesus macaque, a closely related primate. Among the other mammalian species examined, sequence identity to human hGH ranges from 65% to 68%, with sequence similarity ranging from 76% to 79% in rat, mouse, horse, pig, bovine, sheep, and guinea pig. The avian species, common turkey and chicken, show lower sequence identity (55% and 57%, respectively) and similarity (73% and 74%, respectively) to human hGH. The common ostrich exhibits similar values (54% identity and 72% similarity). The lowest sequence identity and similarity are observed in the fish species. Japanese eel shows 44% identity and 61% similarity, goldfish has 38% identity and 58% similarity, and Atlantic salmon displays the lowest values with 36% identity and 52% similarity to human GH.

**Table 1.** Sequence comparison of human growth hormone with growth hormone sequences from other species. The table shows the NCBI and UniProt sequence identifiers, UniProt sequence name, sequence length, percentage of sequence identity, percentage of sequence similarity, and signal peptide length for each species compared to human GH.

| Organism | NCBI Seq ID | Uniprot Seq. ID | Uniprot Seq. Name | Seq length | Seq Identity % | Seq Similarity % | Signal Pep |
| --- | --- | --- | --- | --- | --- | --- | --- |
| Human | NP_000506 | P01241 | SOMA_HUMAN | 217 | 100 | 100 | 26 |
| Rhesus macaque | NP_001036203 | P33093 | SOMA_MACMU | 217 | 96 | 97 | 26 |
| Rat | NP_001030020 | P01244 | SOMA_RAT | 216 | 65 | 76 | 26 |
| Mouse | NP_032143 | P06880 | SOMA_MOUSE | 216 | 67 | 77 | 26 |
| Horse | NP_001075417 | P01245 | SOMA_HORSE | 216 | 67 | 79 | 26 |
| Pig | NP_999034 | P01248 | SOMA_PIG | 216 | 68 | 78 | 26 |
| Bovine | NP_851339 | P01246 | SOMA_BOVIN | 217 | 67 | 77 | 26 |
| Sheep | NP_001009315 | P67930 | SOMA_SHEEP | 217 | 67 | 76 | 26 |
| Guinea pig | NP_001166330 | Q9JKM4 | SOMA_CAVPO | 216 | 65 | 77 | 26 |
| Common turkey | XP_010722827 | P22077 | SOMA_MELGA | 216 | 55 | 73 | 25 |
| Chicken | NP_989690 | P08998 | SOMA_CHICK | 214 | 57 | 74 | 25 |
| Common ostrich | BAA82959 | Q9PWG3 | SOMA_STRCA | 215 | 54 | 72 | 25 |
| Japanese eel | AAA48535 | P08899 | SOMA_ANGJA | 207 | 44 | 61 | 19 |
| Goldfish | AAC19389 | O93359 | SOMA1_CARAU | 210 | 38 | 58 | 22 |
| Atlantic salmon | AAU11454 | Q5SDS1 | Q5SDS1_SALSA | 208 | 36 | 52 | 22 |

The length of the mature growth hormone protein is highly conserved across the mammalian species, ranging from 216 to 217 amino acids, similar to the 217 amino acids in human hGH. The signal peptide length is also conserved at 26 amino acids in most mammals, with a slightly shorter signal peptide of 25 amino acids in the avian species and 19-22 amino acids in the fish species.

#### 3.1.3 Analysis of human GH mutations and their locations in GH protein

Figure 1 provides an overview of the human GH protein sequence and the location of numerous reported mutations. Several key structural features of GH have been shown, including four major α-helices (A, B, C, and D) and two disulfide bonds connecting cysteine residues at positions 53-165 and 182-189. Several point mutations are depicted along the linear sequence, distributed throughout the protein. A significant number of mutations are observed within the α-helical regions, particularly in helix A (residues 9-34) and helix D (residues 158-190), which are known to be critical for binding to the growth hormone receptor (GHR). For instance, mutations like R16C, R16L, R16H, and A17T are located in helix A. Similarly, mutations such as F25Y and F25I are also found in this region. In helix D, mutations like Y164H, K172N, E174K, I179V, I179M, I179S, C182R, R183C, R183H, C189Y, and G190S are present. Mutations are also observed in the loop regions connecting the helices. For example, P2Q, I4T, I4V, R8K, N12H, L15F, H18R, H21Y, A24T, N47K, N47D, L45P are located in the N-terminal region and the loop between helix A and B. Mutations like C53F, C53S, N63K, S62C, R77H, R77C, S79C, Q84E, Q91R, Q91L are found in the loop connecting helix B and C, and within helix C (residues 97-107). In the C-terminal region and the loop between helix C and D, mutations such as T123M, L162P, D169E, D112G, D112H, D116N, D116E, G120C, G120S, and E119D are indicated.

**Figure 1.**
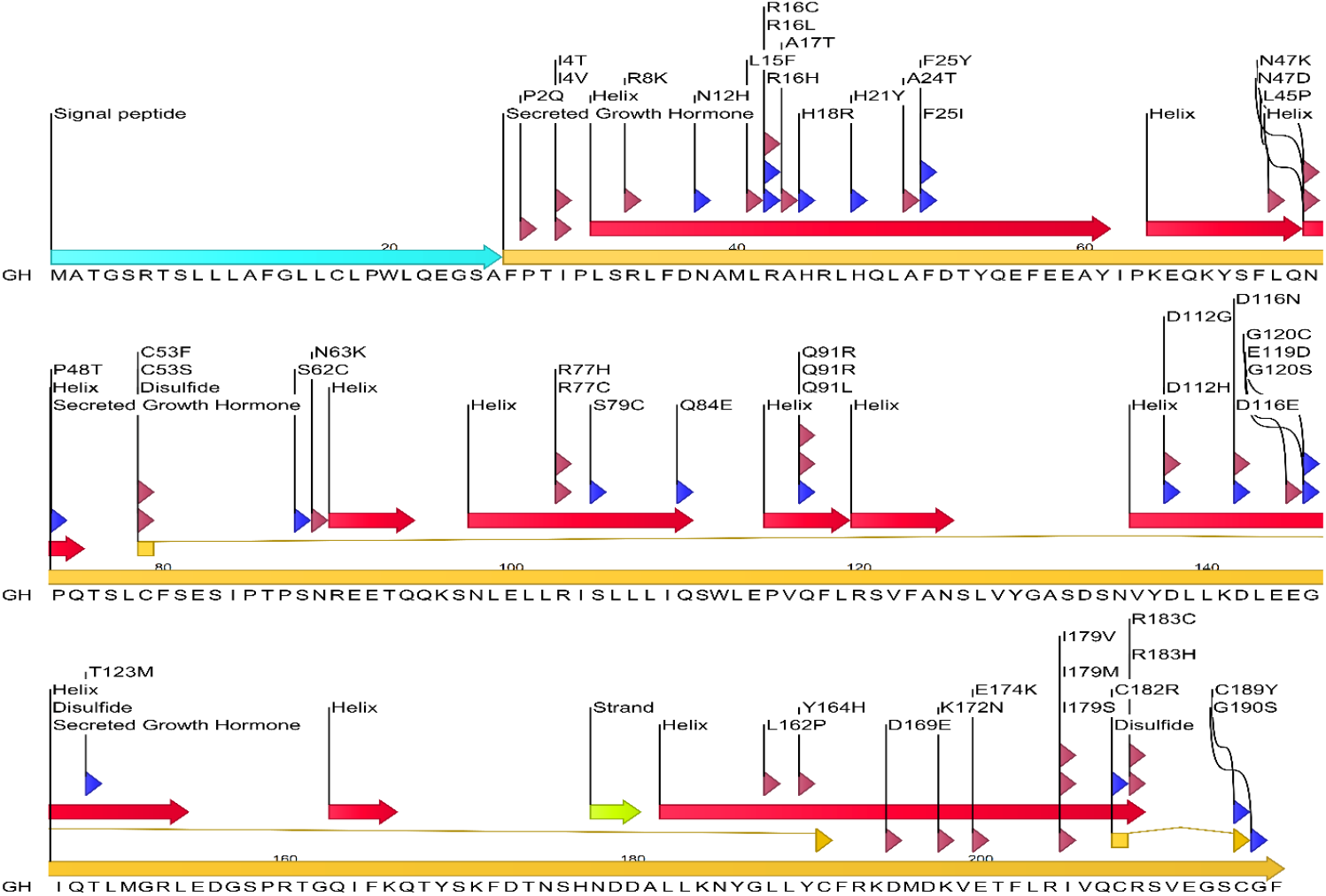
Schematic representation of the human growth hormone protein sequence and location of common mutations. The linear sequence of the secreted 217 amino acid GH protein is shown, indicating key secondary structure elements (α-helices represented by red arrows, β-strand by a yellow arrow, and disulfide bonds by connecting lines). The signal peptide (26 amino acids) is shown at the N-terminus, followed by the mature secreted GH. Various reported mutations are indicated above and below the sequence, with arrows pointing to the affected amino acid residue. Mutations associated with growth hormone deficiency (GHD) or Laron syndrome (LS) are highlighted. The figure illustrates the distribution of these mutations across the GH protein, including within and outside the α-helical regions known to be crucial for interaction with the growth hormone receptor.

Known GH disease-causing mutation data were retrieved from the UniProtKB database (**Table 2**). GH mutations are found to be responsible for two types of diseases: Isolated Growth hormone deficiency (GHD) and Kowarski syndrome (KWKS). The Growth hormone deficiency is an autosomal recessive deficiency of growth hormone leading to short stature. Patients have low but detectable levels of growth hormone, significantly retarded bone age, and a positive response and immunologic tolerance to growth hormone therapy. The Kowarski syndrome is clinically characterized by short stature associated with bio-inactive growth hormone, normal or slightly increased growth hormone secretion, pathologically low insulin-like growth factor 1 levels, and normal catch-up growth on growth hormone replacement therapy. These mutations are distributed throughout the GH protein and exhibit diverse effects on its functional activity. Certain mutations, including L16P and Q117L, predominantly compromise GH secretion. Conversely, mutations such as T53I, K67R, N73D, S97F, and T201A, hinder the capacity of GH to activate the JAK/STAT signaling pathway. Finally, the R209H mutation is associated with autosomal dominant growth hormone deficiency (IGHD2).The HGH variants Thr3, Arg16, Asn47, Gln91, Arg183 and Arg77, Asp112 are associated with growth hormone deficiency [49–51] and Kowarski syndrome [50, 52–54] respectively. It is noteworthy that the R103C mutation, implicated in Kowarski syndrome, does not impede GHR signaling or interaction, but rather demonstrates an augmented interaction with GHBP. The D138G mutation, also associated with KWKS, results in an abrogation of biological activity.

**Table 2.** A List of Disease-Causing Mutations in Human Growth Hormone. The table includes the amino acid change (Original Amino Acid → New Amino Acid), the sequence position in the full-length protein, the associated growth hormone deficiency type or syndrome, the reported effect of the mutation on GH function, the dbSNP identifier (if available), and the relevant publications. Sequence positions might differ depending on whether the signal peptide is included (residues 1-26). The table indicates both the position in the full-length protein and, where available from the source document, the position in the mature secreted growth hormone.

| **Growth hormone deficiency, isolated, 1B (IGHD1B)** |  |  |  |  |  |
| --- | --- | --- | --- | --- | --- |
| **Natural Variant** | **Sequence Position** | **PDB No.** | **Effect of mutation** | **dbSNP** | **Publication** |
| L → P | 16 | - | suppresses secretion |  | [49] Millar |
| D → N | 37 | 11 | - |  | [49] Millar |
| R → C | 42 | 16 | reduced secretion | rs71640273 | [49] Millar |
| T → I | 53 | 27 | reduced ability to activate the JAK/STAT pathway |  | [49] Millar |
| K → R | 67 | 41 | reduced ability to activate the JAK/STAT pathway |  | [49] Millar |
| N → D | 73 | 47 | reduced ability to activate the JAK/STAT pathway | rs71640276 | [49] Millar |
| S → F | 97 | 71 | reduced ability to activate the JAK/STAT pathway |  | [49] Millar |
| E → K | 100 | 74 | - |  | [49] Millar |
| Q → L | 117 | 91 | reduced secretion | Q→R | [49] Millar |
| S → C | 134 | 108 |  |  | [49] Millar |
| S → R | 134 | 108 | reduced ability to activate the JAK/STAT pathway |  | [49] Millar |
| T → A | 201 | 175 | reduced ability to activate the JAK/STAT pathway |  | [49] Millar |
| **Growth hormone deficiency, isolated, 2 (IGHD2)** |  |  |  |  |  |
| R → H | 209 | 183 |  | rs137853223 | [50] Miyata<br>[51] Deladoey |
| **Kowarski syndrome (KWKS)** |  |  |  |  |  |
| R → C | 103 | 77 | No effect on GHR signaling pathway; does not affect interaction with GHR; results in a stronger interaction with GHBP; does not affect the subcellular location. | rs137853220 | [52] Takahashi<br>[53] Petkovic |
| D → G | 138 | 112 | Loss of activity | rs137853221 | [54] Takahashi |

#### 3.1.3 Comparison of human GH protein sequence with diverse GH homologues across species

A multiple sequence alignment (Figure 2) analysis demonstrated a high degree of conservation within the mature growth hormone sequence across a diverse array of vertebrate species, while the N-terminal signal peptide exhibits increased variability. This pattern of conservation and variability is characterized by three principal observations. 1. Extensively Conserved Domains: Discrete segments of amino acids within the mature growth hormone sequence are nearly invariant across mammalian species and demonstrate considerable conservation in more phylogenetically distant species, such as avian and piscine taxa. These extensively conserved domains likely represent critical structural components or functional regions of the growth hormone protein essential for its biological activity. Any alterations within these domains may potentially disrupt the protein’s structural integrity or functional capacity, resulting in deleterious consequences. **2. Invariant Residues:** The alignment also reveals numerous individual amino acid residues that are completely conserved across the majority, if not all, of the aligned sequences. These conserved residues may encompass cysteine residues implicated in disulfide bridge formation, proline residues critical for secondary structure stabilization, or other residues involved in receptor binding or protein-protein interactions. The absolute conservation of these specific residues throughout evolutionary time strongly suggests their indispensable roles in maintaining the protein’s structural framework, stability, receptor affinity, or other essential functions. **3. Variable Segments:** Compared to the highly conserved domains, specific segments within the mature growth hormone sequence exhibit greater variability, particularly when comparing sequences from distantly related species. These variable segments may be involved in species-specific adaptations, facilitating interactions with disparate receptors or responses to divergent physiological conditions. Additionally, these segments may possess greater tolerance to amino acid substitutions without inducing substantial functional impairment.

**Figure 2.**
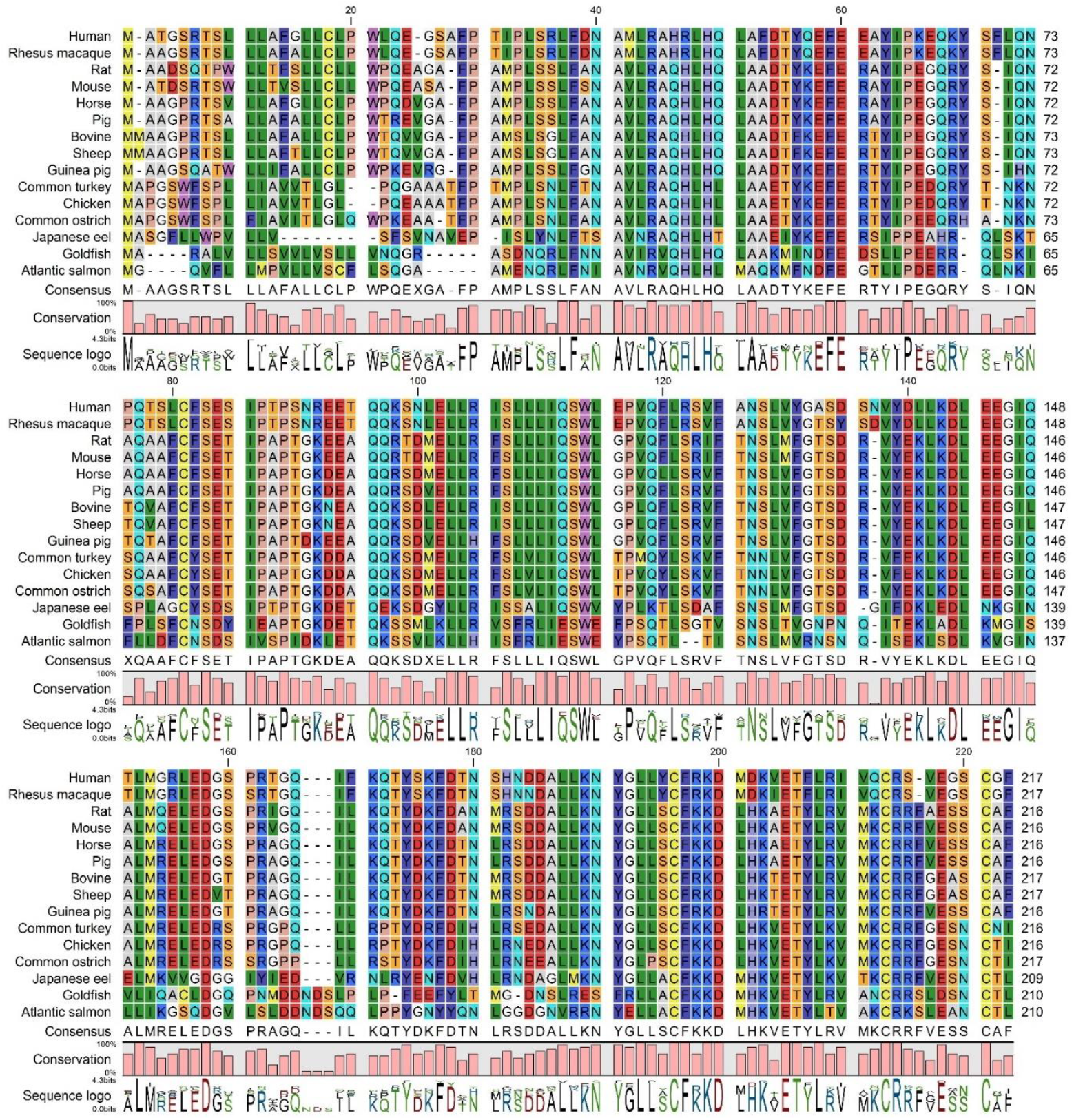
A multiple sequence alignment of human growth hormone alongside growth hormone sequences from various vertebrate species. The alignment encompasses the N-terminal signal peptide, where applicable, and the mature growth hormone sequence. The represented species, arranged from top to bottom, are as follows: Human, Rhesus macaque, Rat, Mouse, Horse, Pig, Bovine, Sheep, Guinea pig, Common turkey, Chicke n, Common ostrich, Japanese eel, Goldfish, and Atlantic salmon. Residues are color-coded based on their chemical characteristics to accentuate patterns of conservation. The consensus sequence and conservation levels are presented beneath the alignment. Notable highly conserved residues and regions are observed throughout the mature growth hormone sequence, indicating their potential functional significance.

### 3.2 Analysis of GH-GHR contacts

#### 3.2.1 Structural analysis of GH-GHR contacts

The growth hormone interacts with its receptor through two distinct binding sites, engaging two receptor molecules. A protein interface contact analysis reveals the specific amino acid residues involved in these interactions.

### Interface 1: GH (Chain A) - GHR (Chain B)

Key hydrogen bonds are observed between the following residue pairs (hGH residue - GHR residue): Phe25-Ser219, Lys41-Met170, Arg167-Glu127, Tyr164-Gly220, Arg178-Glu127, Gly190-Ile165, Cys189-Gln166, Leu45-Trp76, Tyr42-Lys121, Pro48-Asn218, Glu174-Lys167, His21-Glu44, His18-Arg217, Gln22-Arg217, Ile179-Ser102, Cys182-Gly168, Lys172-Phe123, Ser62-Asp164, Arg64-Asp164, Gln46-Trp169, Asn63-Glu44, Thr175-Lys167, Pro61-Thr77, Asp171-Arg43, and Lys168-Trp104. There are extensive hydrophobic interactions at the interface, involving the same residue pairs listed above for hydrogen bonds, along with additional contacts. Notably, residues such as Phe25, Lys41, Arg167, Tyr164, Gly190, Cys189, Leu45, Tyr42, Pro48, Glu174, His21, His18, Gln22, Ile179, Cys182, Lys172, Ser62, Arg64, Gln46, Asn63, Thr175, Pro61, Asp171, and Lys168 in hGH make hydrophobic contacts with residues in the GHR.

### Interface 2: GH (Chain A) - GHR (Chain C)

Analysis of this interface reveals a separate set of interactions including hydrogen Bonds: Leu9 (A) - Asp126 (C), Thr123 (A) - Trp104 (C), Gly120 (A) - Ser102 (C), Asp116 (A) - Trp104 (C), Glu119 (A) - Ser102 (C), Arg16 (A) - Glu44 (C), Ile4 (A) - Ile103 (C), Leu15 (A) - Gly168 (C), Asn12 (A) - Arg43 (C), Arg8 (A) - Asp126 (C), Tyr103 (A) - Ile165 (C), Pro2 (A) Pro106 (C), and Phe1 (A) - Arg71 (C). Multiple hydrophobic Contacts between the two chains, similar to interactions seen between GH and GHR chain B, notably: Leu9 (A) - Asp126 (C), Thr123 (A) - Trp104 (C), Gly120 (A) - Ser102 (C), Asp116 (A) - Trp104 (C), Glu119 (A) - Ser102 (C), Arg16 (A) - Glu44 (C), Ile4 (A) - Ile103 (C), Leu15 (A) - Gly168 (C), Asn12 (A) - Arg43 (C), Arg8 (A) - Asp126 (C), Tyr103 (A) - Ile165 (C), Pro2 (A) - Pro106 (C), and Phe1 (A) - Arg71 (C).

Mapping of Common Mutations to the Interface: Comparison with the locations of common GH mutations (Figure 1) reveals that several reported mutations occur at residues directly involved in the hGH-GHR interface:

- His18: The mutation H18R is at an interface residue interacting with Arg217 of the GHR.
- His21: The mutation H21Y is at an interface residue interacting with Glu44 of the GHR.
- Phe25: Mutations F25Y and F25I are located at a key interface residue forming both hydrogen bonds and hydrophobic contacts with Ser219 of the GHR.
- Lys41: Lys41 forms interactions with Met170 of the GHR.
- Leu45: The mutation L45P is at an interface residue interacting with Trp76 of the GHR.
- Pro48: The mutation P48T is located at a residue involved in interactions with Asn218 of the GHR.
- Ile179: Mutations I179V, I179M, and I179S are located at a residue interacting with Ser102 of the GHR.
- Cys182: The mutation C182R is at an interface residue forming interactions with Gly168 of the GHR.
- Lys172: The mutation K172N is at a residue interacting with Phe123 of the GHR.
- Ser62: The mutation S62C is at an interface residue interacting with Asp164 of the GHR.
- Asn63: The mutation N63K is at an interface residue interacting with Glu44 of the GHR.
- Pro61: Pro61 interacts with Thr77 of the GHR.
- Lys168: Lys168 interacts with Trp104 of the GHR.
- Tyr164: The mutation Y164H is at a key interface residue interacting with Gly220 of the GHR.
- Glu174: The mutation E174K is at a residue interacting with Lys167 of the GHR.
- Gly190: The mutation G190S is at an interface residue interacting with Ile165 of the GHR.
- Cys189: The mutation C189Y is at an interface residue interacting with Gln166 of the GHR.
- Arg16: Mutations R16C, R16L, and R16H are located in close proximity to the interface and might indirectly affect binding.

Mapping of Common Mutations to the Second Interface: Several reported mutations occur at residues involved in this second hGH-GHR interface:

- Phe1: Phe1 interacts with Arg71 of GHR chain C.
- Pro2: The mutation P2Q is located at an interface residue interacting with Pro106 of GHR chain C.
- Ile4: Mutations I4T and I4V are at interface residues interacting with Ile103 of GHR chain C.
- Arg8: The mutation R8K is located at a residue interacting with Asp126 of GHR chain C.
- Asn12: The mutation N12H is at an interface residue interacting with Arg43 of GHR chain C.
- Leu15: The mutation L15F is at an interface residue interacting with Gly168 of GHR chain C.
- Arg16: Mutations R16C, R16L, and R16H are located at a key interface residue interacting with Glu44 of GHR chain C.
- Leu9: Leu9 interacts with Asp126 of GHR chain C.
- Gly120: Mutations G120C and G120S are at interface residues interacting with Ser102 of GHR chain C.
- Asp116: Mutations D116N and D116E are at interface residues interacting with Trp104 of GHR chain C.
- Glu119: The mutation E119D is at an interface residue interacting with Ser102 of GHR chain C.
- Thr123: The mutation T123M is at an interface residue interacting with Trp104 of GHR chain C.
- Tyr103: Tyr103 interacts with Ile165 of GHR chain C.

There are overlaps in the interaction of the two GHR chains (B and C) with GH (chain A). Many residues on GH (chain A) appear to interact with both chain B and chain C of the GHR:

- Arg16 forms a hydrogen bond with Glu44 on chain C and is in close proximity to the interface with chain B
- Gly120 forms a hydrogen bond with Ser102 on chain C and is likely involved in interactions within the first binding site.
- Asp116 forms a hydrogen bond with Trp104 on chain C and is likely involved in interactions within the first binding site.
- Glu119 forms a hydrogen bond with Ser102 on chain C and is likely involved in interactions within the first binding site.
- Thr123 forms a hydrogen bond and hydrophobic contact with Trp104 on chain C and is likely involved in interactions within the first binding site.

These overlaps highlight the complex nature of the hGH interaction with its dimeric receptor, where certain regions of GH are crucial for engaging both receptor chains to facilitate dimerization and subsequent signaling.

Comparison with our contact analysis (Figure 3 and 4) reveals a significant overlap in the identified contact residues for both GHR chains. Most of the GH residues listed as contacting either chain B or chain C in this table were also highlighted in the respective contact figures, validating the consistency of the contact mapping.

**Figure 3.**
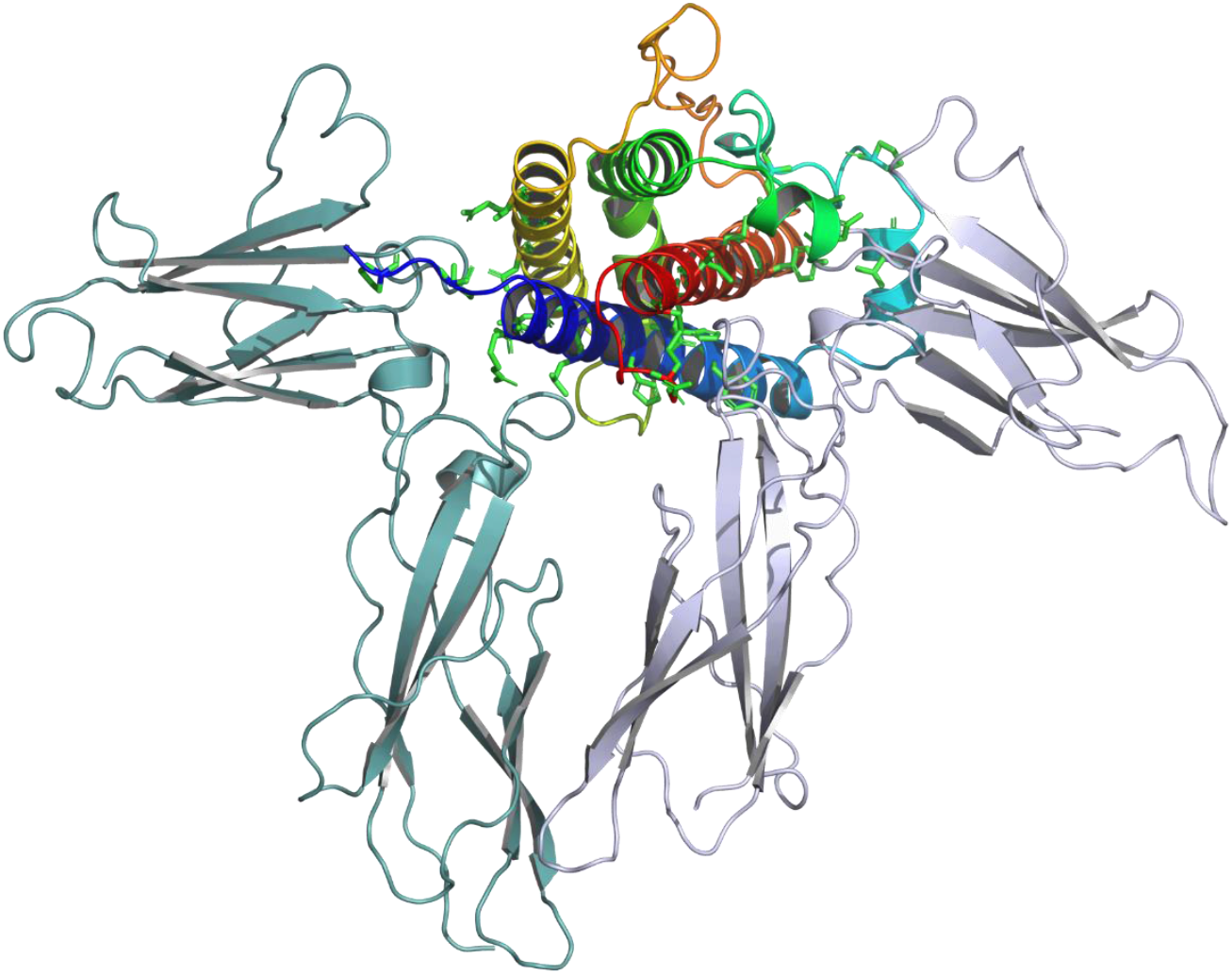
Crystal Structure of the Human Growth Hormone in Complex with Two Extracellular Domains of the Growth Hormone Receptor (GHR) (PDB ID: 3HHR). hGH, Chain A, is presented as a rainbow ribbon diagram. The two GHR molecules, denoted as Receptor 1 and Receptor 2 to indicate their sequential binding order, are represented in light green and light blue, respectively. These extracellular GHR domains mediate the recognition and binding of GH. The structural arrangement demonstrates the sequential binding model of GHR dimerization. Initially, GH engages with Receptor 1 through Site 1, inducing conformational alterations in GH that expose Site 2 for subsequent interaction with Receptor 2. The formation of a GHR dimer occurs upon the binding of two GHR molecules to a single hGH molecule. Dimerization of GHR is imperative for its activation. The dimerized receptor complex initiates intracellular signaling cascades, including the JAK-STAT pathway, resulting in diverse physiological effects such as cellular growth, proliferation, and differentiation.

**Figure 4.**
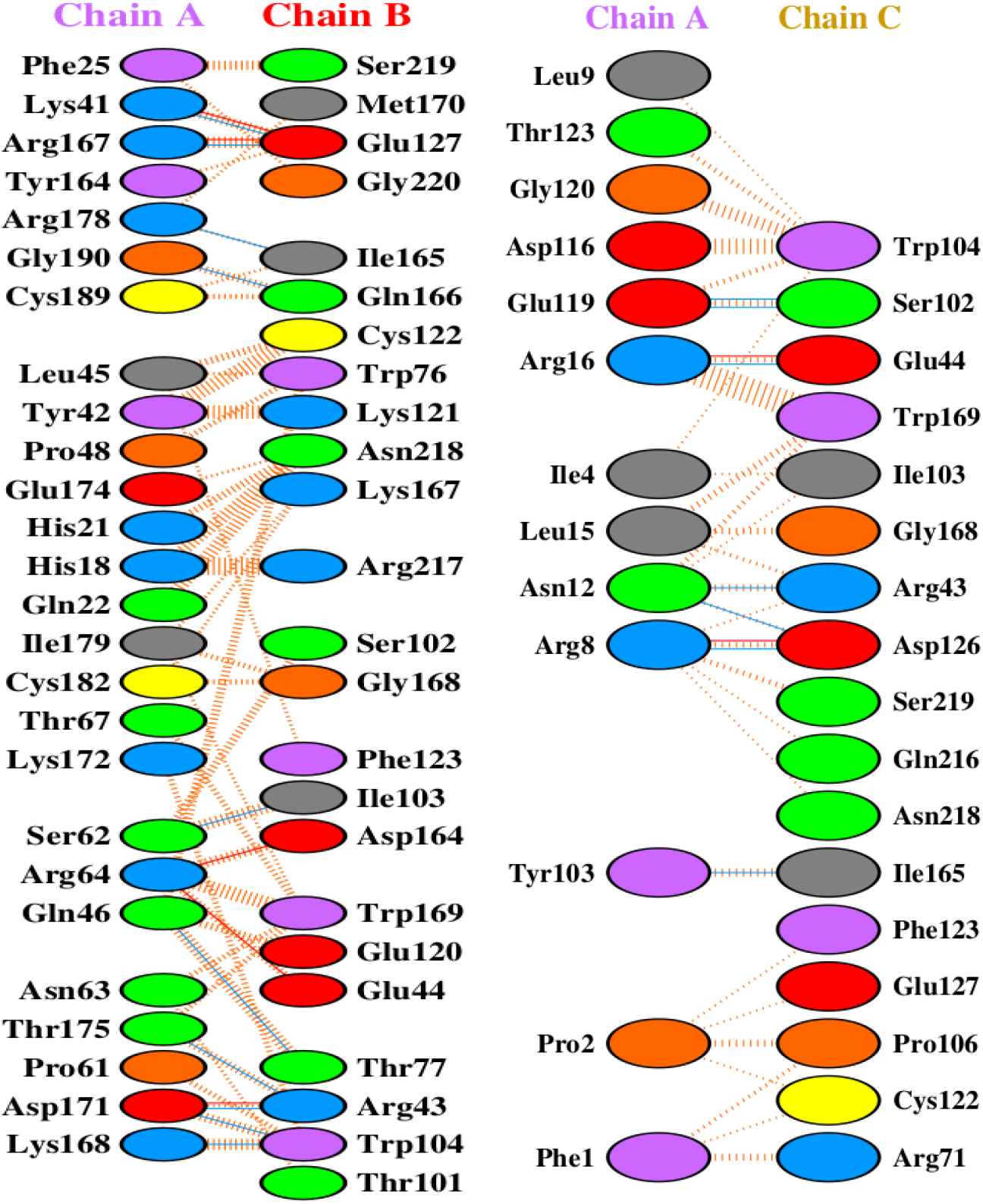
Contact analysis of the two binding interfaces between human growth hormone (chain A) and its receptor (chains B and C). Panel A shows the interactions between hGH (chain A) and one receptor chain (chain B). Panel B shows the interactions between hGH (chain A) and the second receptor chain (chain C). In both panels, hydrogen bonds are depicted as dashed orange lines, and hydrophobic contacts as arcs with spokes. The figures illustrate the residues in hGH crucial for engaging both receptor chains in the functional complex.

### 3.3 Analysis of sequence conservation pattern in growth hormone at GH-GHR contact points

Analysis of the ConSurf conservation scores for these contact residues shows a general trend towards higher conservation. Many of the GH residues involved in receptor binding exhibit scores of 6 or higher, indicating their evolutionary importance (Figure 5). For instance, residues such as R16, H18, H21, F25, Y42, Y47, F51, L45, P48, S62, N63, Q91, Y103, D116, E119, T123, Y164, R167, K172, E174, R178, I179, C182, C189, G190, and F1, P2, I4, R8, N12, L15, L9, G120, and T123 (contacting chain C) display notable conservation. Furthermore, several GH residues that are known to be mutated in human diseases (UniProt Disease Variant column) are also present in table 2. Many of these clinically significant residues, such as P2, I4, R8, N12, L15, R16 (R>C), A17, H18, H21, A24, F25, L45, N47 (N>D), P48, S62, N63, Q91, D112, D116, E119, G120, T123, L162, Y164, D169, K172, E174, I179, C182, R183, C189, and G190, are also identified as contact residues for one or both chains of the GHR.

**Figure 5.**
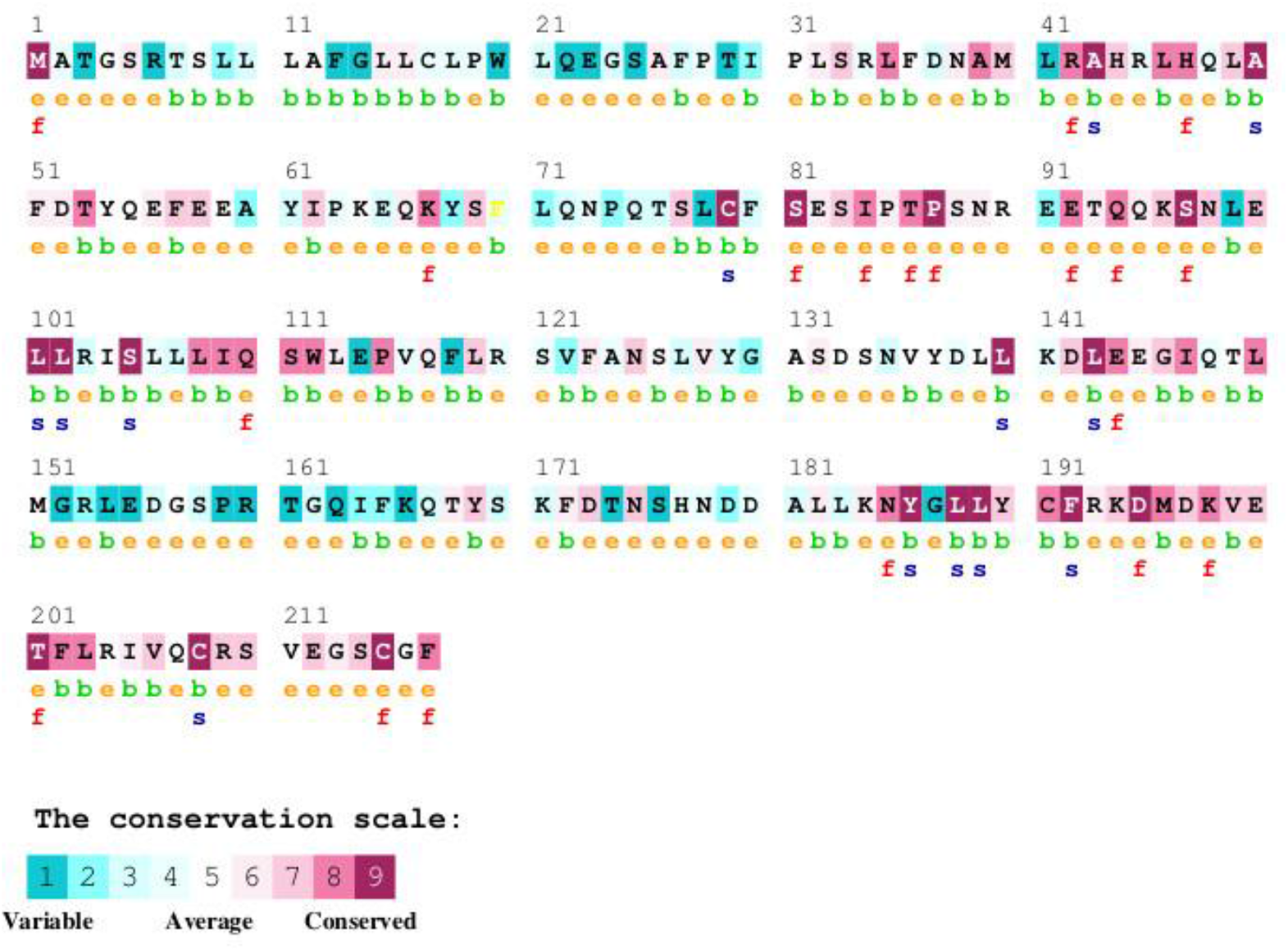
ConSurf analysis of the human growth hormone (hGH) protein showing sequence conservation. The amino acid sequence of hGH is displayed, with each residue colored according to its conservation level, ranging from variable (cyan to blue) through average (green) to conserved (yellow to maroon). The conservation scores are based on a multiple sequence alignment of hGH orthologs from various species. Predicted functional residues (highly conserved and exposed) are marked with an ‘f’, and predicted structural residues (highly conserved and buried) are marked with an ‘s’. Exposed residues are also indicated with an ‘e’, and buried residues with a ‘b’. Residues with insufficient data for conservation calculation are marked with an ‘X’.

The ConSurf analysis (Figure 5, Table 4) reveals a significant degree of sequence conservation across the human growth hormone protein, with several regions exhibiting high conservation scores. The N-terminal region of the mature hGH (residues 1-~40) shows a mix of conservation levels, with some highly conserved patches interspersed with more variable residues. Notably, residues R16, H18, H21, and F25 within this region, which are known to be involved in receptor binding (as shown in Figure 3 and 4), display high conservation. The region encompassing helix C and the loop connecting it to helix D (approximately residues 95-145) also shows considerable conservation, with several residues predicted to be functional (‘f’) or structural (‘s’). Specifically, residues like Q91, Y103, and D116, which participate in receptor binding, are located within this conserved segment (Figure 2). The C-terminal helix D (residues ~158-190), another crucial region for receptor interaction, exhibits a high degree of conservation. Residues Y164, R167, K172, E174, R178, I179, C182, C189, and G190, all implicated in binding to the growth hormone receptor (Figure 2), are located within this highly conserved helix. Several of these are also predicted to be functional (‘f’) or structural (‘s’). In contrast, some loop regions and the very C-terminus (residues 191-217) appear to be more variable, suggesting less stringent evolutionary constraints on these parts of the protein.

**Table 3.**
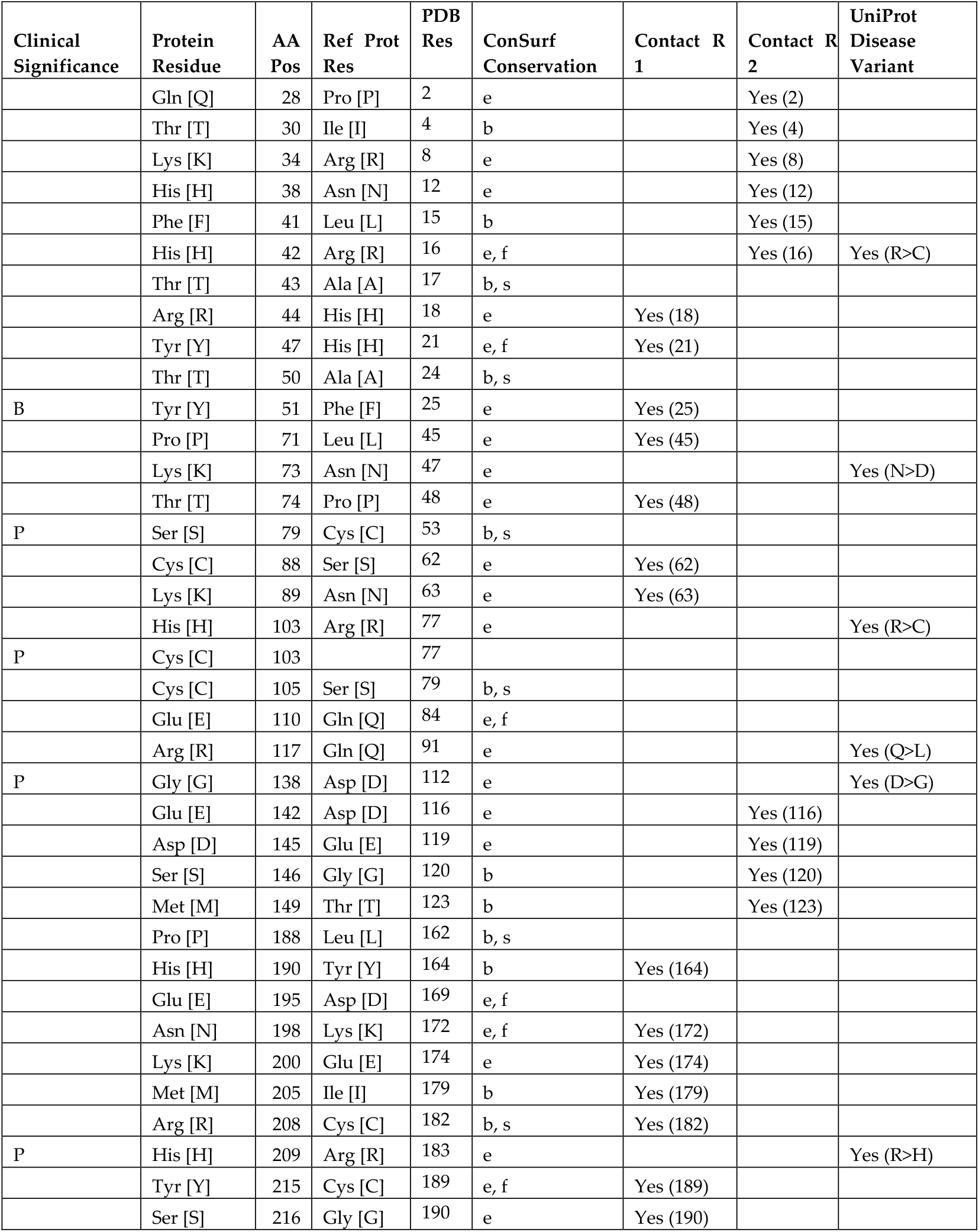
A summary of amino acid contacts between human growth hormone (GH) and its receptor (GHR) (chains B and C). The table lists the GH residue (Mutated Protein Residue), its position (AA Pos), the corresponding residue in GHR chain B (Ref Prot Residue), the PDB residue number, the ConSurf conservation score of the GH residue, whether it was identified as a contact residue in LigPlot+ analysis for chain B (Contact R 1) and chain C (Contact R 2), and if it is a known disease variant according to UniProt (UniProt Disease Variant).

| Clinical Significance | Protein Residue | AA Pos | Ref Prot Res | PDB Res | ConSurf Conservation | Contact R 1 | Contact R 2 | UniProt Disease Variant |
| --- | --- | --- | --- | --- | --- | --- | --- | --- |
|  | Gln [Q] | 28 | Pro [P] | 2 | e |  | Yes (2) |  |
|  | Thr [T] | 30 | Ile [I] | 4 | b |  | Yes (4) |  |
|  | Lys [K] | 34 | Arg [R] | 8 | e |  | Yes (8) |  |
|  | His [H] | 38 | Asn [N] | 12 | e |  | Yes (12) |  |
|  | Phe [F] | 41 | Leu [L] | 15 | b |  | Yes (15) |  |
|  | His [H] | 42 | Arg [R] | 16 | e, f |  | Yes (16) | Yes (R>C) |
|  | Thr [T] | 43 | Ala [A] | 17 | b, s |  |  |  |
|  | Arg [R] | 44 | His [H] | 18 | e | Yes (18) |  |  |
|  | Tyr [Y] | 47 | His [H] | 21 | e, f | Yes (21) |  |  |
|  | Thr [T] | 50 | Ala [A] | 24 | b, s |  |  |  |
| B | Tyr [Y] | 51 | Phe [F] | 25 | e | Yes (25) |  |  |
|  | Pro [P] | 71 | Leu [L] | 45 | e | Yes (45) |  |  |
|  | Lys [K] | 73 | Asn [N] | 47 | e |  |  | Yes (N>D) |
|  | Thr [T] | 74 | Pro [P] | 48 | e | Yes (48) |  |  |
| P | Ser [S] | 79 | Cys [C] | 53 | b, s |  |  |  |
|  | Cys [C] | 88 | Ser [S] | 62 | e | Yes (62) |  |  |
|  | Lys [K] | 89 | Asn [N] | 63 | e | Yes (63) |  |  |
|  | His [H] | 103 | Arg [R] | 77 | e |  |  | Yes (R>C) |
| P | Cys [C] | 103 |  | 77 |  |  |  |  |
|  | Cys [C] | 105 | Ser [S] | 79 | b, s |  |  |  |
|  | Glu [E] | 110 | Gln [Q] | 84 | e, f |  |  |  |
|  | Arg [R] | 117 | Gln [Q] | 91 | e |  |  | Yes (Q>L) |
| P | Gly [G] | 138 | Asp [D] | 112 | e |  |  | Yes (D>G) |
|  | Glu [E] | 142 | Asp [D] | 116 | e |  | Yes (116) |  |
|  | Asp [D] | 145 | Glu [E] | 119 | e |  | Yes (119) |  |
|  | Ser [S] | 146 | Gly [G] | 120 | b |  | Yes (120) |  |
|  | Met [M] | 149 | Thr [T] | 123 | b |  | Yes (123) |  |
|  | Pro [P] | 188 | Leu [L] | 162 | b, s |  |  |  |
|  | His [H] | 190 | Tyr [Y] | 164 | b | Yes (164) |  |  |
|  | Glu [E] | 195 | Asp [D] | 169 | e, f |  |  |  |
|  | Asn [N] | 198 | Lys [K] | 172 | e, f | Yes (172) |  |  |
|  | Lys [K] | 200 | Glu [E] | 174 | e | Yes (174) |  |  |
|  | Met [M] | 205 | Ile [I] | 179 | b | Yes (179) |  |  |
|  | Arg [R] | 208 | Cys [C] | 182 | b, s | Yes (182) |  |  |
| P | His [H] | 209 | Arg [R] | 183 | e |  |  | Yes (R>H) |
|  | Tyr [Y] | 215 | Cys [C] | 189 | e, f | Yes (189) |  |  |
|  | Ser [S] | 216 | Gly [G] | 190 | e | Yes (190) |  |  |

**Table 4.** ConSurf amino acid conservation scores for selected mutations in the human growth hormone protein. Conservation scores range from 1 (variable) to 9 (highly conserved). The residue variety across species indicates the different amino acids observed at that position in orthologous sequences.

| SNV | Conservation score | Residue variety across species |
| --- | --- | --- |
| P2Q | 8 | P,Y,V |
| I4T | 5 | A,F,T,P,E,V,M,I,L |
| R8K | 4 | S,W,N,K,E,H,Q,D,R,G |
| N12H | 7 | S,T,N,K,E,H,M,C,I,R,L |
| L15F | 1 | S,F,T,N,K,E,V,H,Q,M,R,I,G,L |
| R16H, R16L, R16C | 7 | H,Q,R,Y,L,V |
| A17T | 8 | S,A,T,I,L,V |
| H18R | 7 | S,W,T,N,E,H,Q,D |
| H21Y | 7 | F,S,H,K,R,Y,V |
| A24T | 8 | S,A,T,N,Y,V |
| F25Y, F25I | 6 | S,A,F,T,K,E,Y,Q,M,D,R,I,G,L |
| L45P | 6 | L |
| N47K, N47D | 3 | S,A,T,N,P,K,V,H,M,D,I,G |
| C53S, C53F | 9 | C |
| S62C | 5 | A,S,T,N,K,E,V,H,Q,M,I,G |
| N63K | 7 | S,D,N,P,G,E |
| R77H, R77C | 5 | S,N,K,H,Q,D,R,G,L |
| S79C | 7 | A,S,M,T,I,G,V |
| Q84F | 6 | S,W,P,Y,E,V,H,Q,M,D,R,I,L |
| Q91R | 3 | S,F,A,N,K,E,Y,V,H,Q,D,R,I,G,L |
| D112G, G112H | 1 | A,S,T,N,K,P,E,H,Q,D,R,G,L |
| D116E, D116N | 4 | A,S,N,K,E,Y,V,Q,D,R,I,G |
| E119D | 4 | S,A,T,N,K,E,V,Q,M,D,R,L |
| G120S, G120C | 9 | A,F,T,G,Y |
| T123M | 2 | S,A,T,N,K,E,V,M,R,I,L |
| L162P | 7 | F,T,M,N,K,I,L,V |
| Y164H | 6 | A,S,T,N,Y,H,M,C,R |
| D169E | 9 | D,E |
| K172N | 9 | H,M,N,R,K |
| E174K | 9 | S,Q,D,Y,E |
| I179V, I179M, I179S | 7 | F,T,M,I,L,V |
| C182R | 9 | C |
| R183H, R183C | 9 | Q,K,R |
| C189Y | 9 | C |
| G190S | 4 | S,A,T,G |

ConSurf conservation scores for a selected list of mutations in the human growth hormone protein were analyzed (Table 4). The conservation scores range from 1 to 9, with higher scores indicating greater evolutionary conservation. Several disease-associated mutations are found in residues with high conservation scores. For instance, P2Q and A17T both have a conservation score of 8, while N12H and “R16H, R16L, R16C” have a score of 7. This suggests that these positions are important for the protein’s function or structure, as they have been conserved across evolution. Mutations at these highly conserved sites might therefore disrupt normal protein activity, leading to disease. Conversely, some mutations occur at positions with lower conservation scores. L15F has a conservation score of 1, and “D112G, G112H” has a score of 1, indicating these residues are more variable across species. Mutations at these positions might have less impact on the protein’s overall function, although they are still associated with disease. The residue variety across species further supports the conservation scores. For example, C53S and C53F have a conservation score of 9, and the residue variety shows only C at this position across species, highlighting its critical role. In contrast, L15F has a conservation score of 1, with a high variety of amino acids (S, F, T, N, K, E, V, H, Q, M, R, I, G, L) observed at this position in other species, suggesting it is less constrained.

### 3.4 Stability analysis of GH mutations

All selected HGH variants from previous analyses were subjected to site-directed mutagenesis via the SDM tool (Table 5). SDM software predicts the effect of the mutation on protein stability and potentiality as a disease-causing mutant. SDM uses a set of conformationally-constrained environment-specific scoring matrices to calculate the difference in stability (pseudo ΔΔG score) between the wild-type and mutant protein upon providing a structure of wild-type and a list of mutations [43, 44]. The negative and positive values of pseudo ΔΔG correspond to mutations predicted to be destabilizing and stabilizing, respectively. Besides the pseudo ΔΔG score SDM output also provide information about various structural features, including class of secondary structure (SSE), solvent accessibility (RSA), residue depth (DEPT), occluded surface packing (OSP) sidechain-sidechain hydrogen bonding (SS), sidechain-mainchain amide hydrogen bonding (SN) and sidechain-mainchain carbonyl hydrogen bonding (SO) for the wild-type and mutant residues [44]. Previously, it has been predicted that the highly destabilizing mutations are mostly found at high residue packing density regions (OSP > 0.56) and occur at two distinct depth levels (4 Å and 8 Å) and highly stabilizing mutations were observed to occur mostly at high packing density regions and residue depth ~4 Å [44].

**Table 5.**
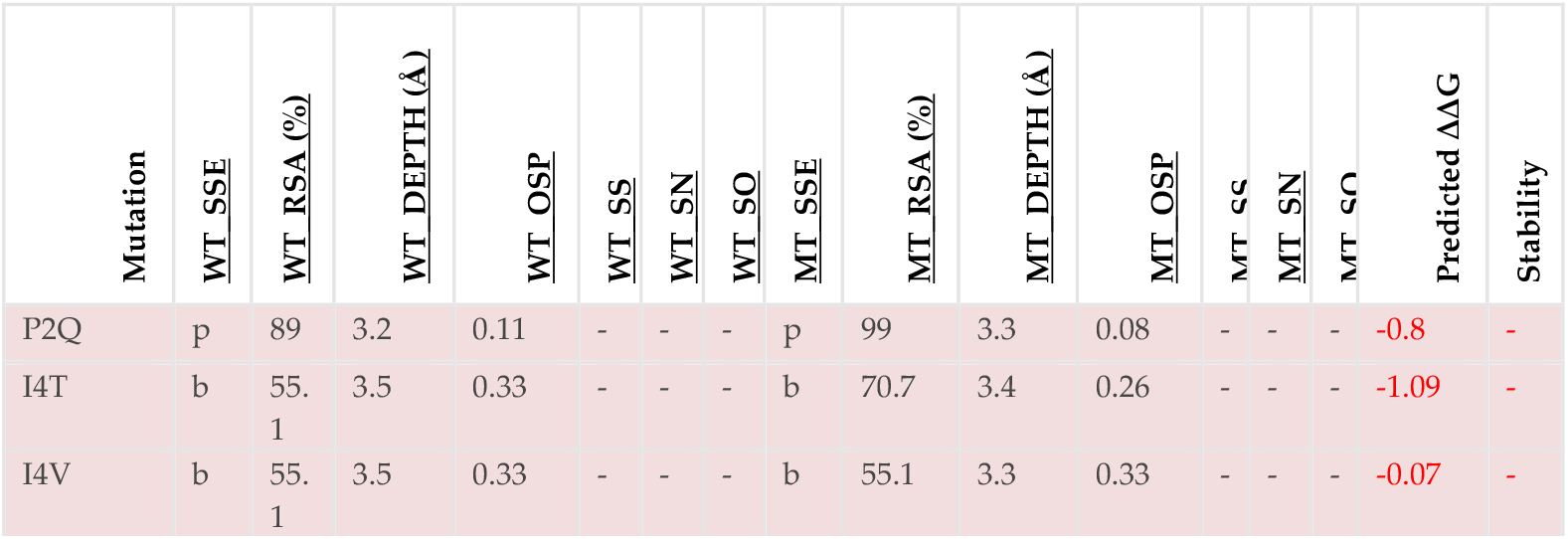
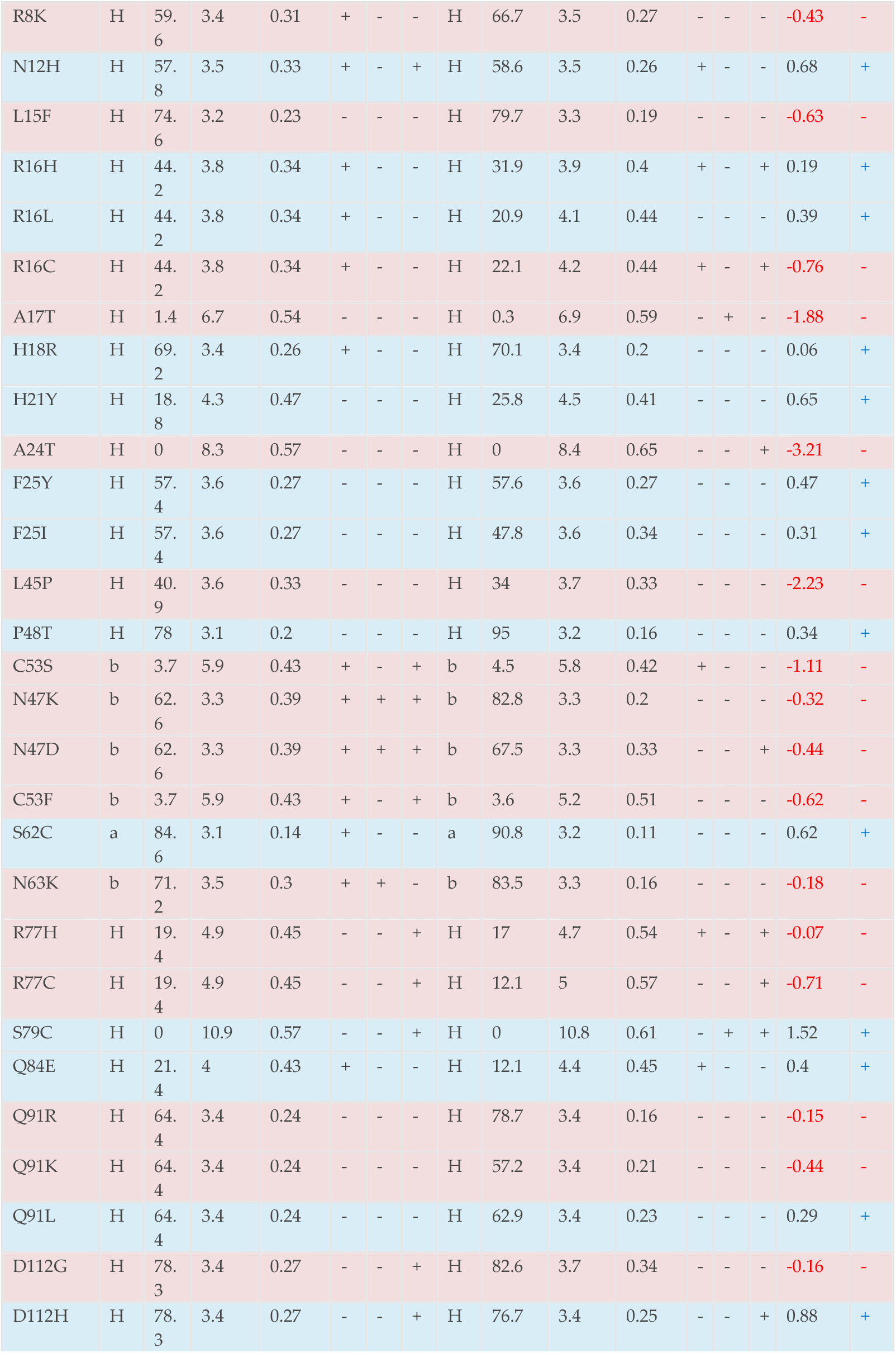
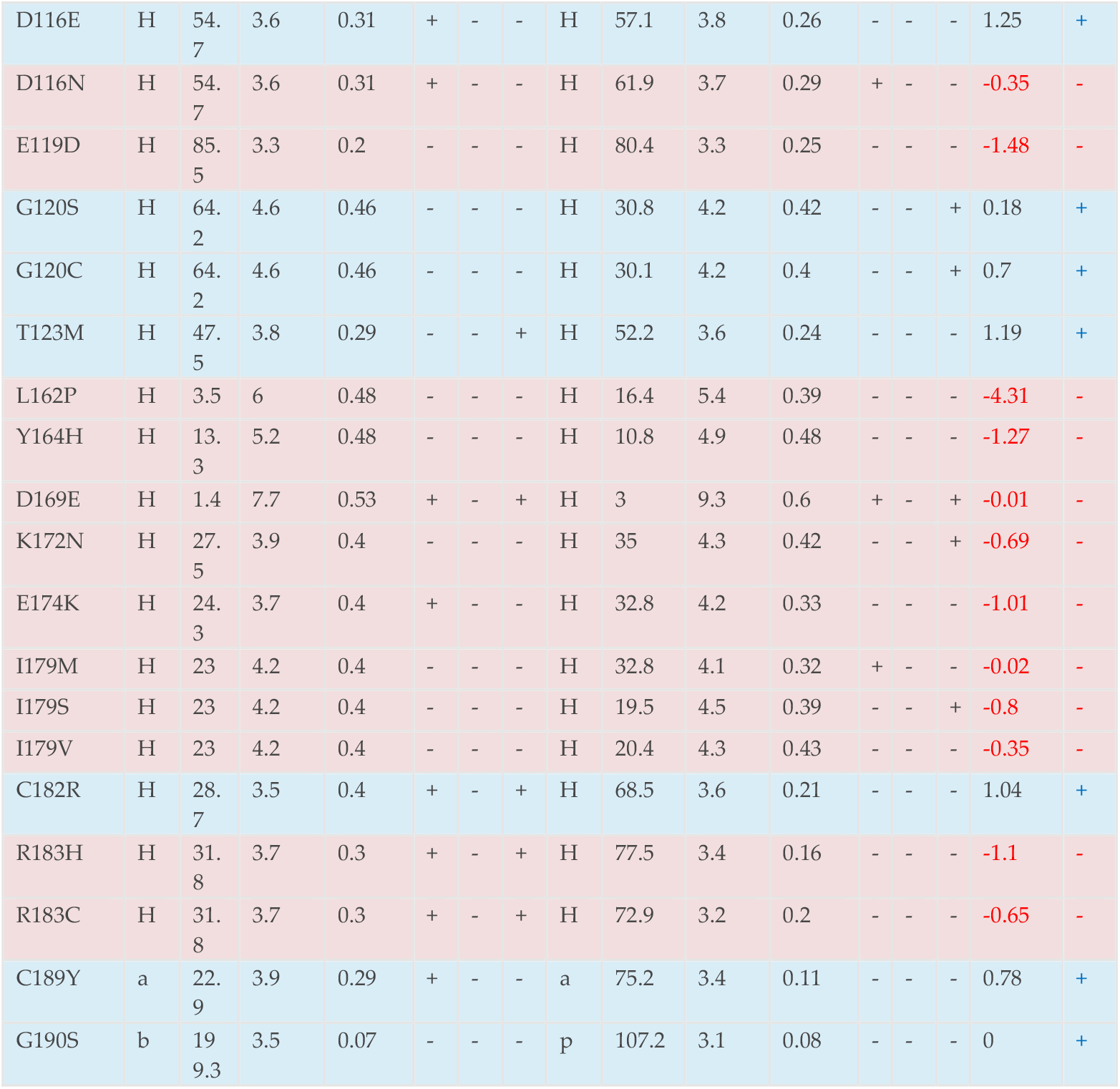
Impact of amino acid changes caused by human genetic variations on GH residues involved in contact with GHR. Amino acids changes reported at positions identified as involved in binding to GHR were collected from NCBI genomic database. After making a contact map of GH-GHR interactions, all amino acid variations reported in GH1 gene were searched and variations occurring at amino acids of GH making contact with GHR were further analyzed by virtual site direct mutagenesis tool (SDM). Predicted changes in protein stability (ΔΔG) selected mutations in the human growth hormone protein, as analyzed by the SDM tool. The table includes the mutated residue (Mutation), secondary structure element (SSE) and relative solvent accessibility (RSA) for both the wild-type (WT) and mutant (MT) proteins, residue depth (DEPTH), outer shell potential (OSP), structural class (SS), structural neighborhood (SN), and solvent organization (SO). The predicted ΔΔG (kcal/mol) indicates the change in protein stability upon mutation, with a positive value suggesting decreased stability and a negative value suggesting increased stability. The Stability column indicates whether the mutation is predicted to destabilize (−) or stabilize (+) the protein.

In our analysis, we have found that out of 51 HGH variants 31 are responsible for the reduced stability of the proteinThe Site-Directed Mutagenesis (SDM) analysis predicted a range of stability changes (ΔΔG) for the selected mutations in the human growth hormone protein (Table 5). The predicted ΔΔG values ranged from −4.31 kcal/mol to 1.52 kcal/mol, indicating that some mutations are predicted to increase protein stability while others are predicted to decrease it.

Several mutations were predicted to significantly destabilize the protein (negative ΔΔG values). These include L162P (−4.31 kcal/mol), A24T (−3.21 kcal/mol), L45P (−2.23 kcal/mol), A17T (−1.88 kcal/mol), E119D (−1.48 kcal/mol), Y164H (−1.27 kcal/mol), I4T (−1.09 kcal/mol), C53S (−1.11 kcal/mol), R183H (−1.1 kcal/mol), and E174K (−1.01 kcal/mol). Conversely, some mutations were predicted to increase protein stability (positive ΔΔG values), such as S79C (1.52 kcal/mol), D116E (1.25 kcal/mol), T123M (1.19 kcal/mol), C182R (1.04 kcal/mol), D112H (0.88 kcal/mol), C189Y (0.78 kcal/mol), G120C (0.7 kcal/mol), N12H (0.68 kcal/mol), H21Y (0.65 kcal/mol), S62C (0.62 kcal/mol), F25Y (0.47 kcal/mol), Q84E (0.4 kcal/mol), R16L (0.39 kcal/mol), P48T (0.34 kcal/mol), F25I (0.31 kcal/mol), Q91L (0.29 kcal/mol), R16H (0.19 kcal/mol), G120S (0.18 kcal/mol), and H18R (0.06 kcal/mol).

Other mutations showed minimal predicted changes in stability, with ΔΔG values close to zero, such as I4V (−0.07 kcal/mol), R77H (−0.07 kcal/mol), D169E (−0.01 kcal/mol), and I179M (−0.02 kcal/mol). The HGH variants L162P and A17T scored higher pseudo ΔΔG values (−4.31, −3.21 respectively) and are linked with growth hormone deficiency. Similarly, variant C53S (ΔΔG −1.11) is linked with Kowarski syndrome.

## 4. Discussion

The sequence comparison of human growth hormone with its orthologs across a diverse range of species highlights the evolutionary conservation of this important hormone. The high degree of sequence identity and similarity observed in primates underscores the close evolutionary relationship and likely conservation of function. The moderate sequence identity and similarity in other mammalian species suggest a conserved core function of growth hormone, although some species-specific variations might exist, potentially reflecting adaptations to different physiological needs or receptor interactions. The lower sequence identity and similarity in avian and fish species indicate a greater evolutionary distance, which is expected given the phylogenetic relationships. However, the fact that significant sequence similarity is still present suggests that the fundamental roles of growth hormone have been maintained throughout vertebrate evolution.

Mutations within the highly conserved regions of the human growth hormone gene are more probable to impair the normal protein function. Such mutations may precipitate various growth disorders, including growth hormone deficiency or Laron syndrome. The specific conserved residues identified warrant further investigation, as mutations at these sites in human GH may have severe functional consequences, potentially affecting protein folding, secretion, receptor binding affinity, or downstream signaling. These findings align with the ConSurf analysis, where highly conserved regions in human growth hormone are likely to correspond to residues that are identical or highly similar in these other species. These conserved residues are likely crucial for maintaining the protein’s structure, stability, and interactions with the growth hormone receptor.The variations in sequence identity and similarity across species can also provide insights into which regions of the protein might be more amenable to change and potentially responsible for species-specific effects of growth hormone.

The spectrum of GH1 mutations leading to growth disorders underscores the critical importance of various structural and functional aspects of the growth hormone protein. Mutations affecting secretion likely disrupt the processes involved in GH synthesis, post-translational modification, or release from somatotroph cells in the anterior pituitary. Impairment of JAK/STAT pathway activation indicates that these mutations might affect the interaction of GH with its receptor (GHR) or the subsequent conformational changes required for downstream signaling. The unique case of the R103C mutation in Kowarski syndrome, which enhances GHBP interaction without affecting GHR binding or signaling, suggests a more complex regulatory role for GHBP in GH action or availability. The D138G mutation leading to a complete loss of activity likely results in a severely misfolded or non-functional GH protein. The identification of these and other mutations provides valuable insights into the genotype-phenotype correlations in growth hormone deficiency and related syndromes.

The clustering of mutations within the α-helical regions, particularly helix A and D, which are known to form the primary binding interface with the GHR, suggests that these mutations are highly likely to disrupt the crucial contacts required for receptor activation. Amino acid substitutions in these regions can alter the shape, charge, or hydrophobicity of the binding surface, potentially leading to reduced binding affinity or complete loss of interaction, as observed in some forms of growth hormone deficiency (GHD) and Laron syndrome (LS). Mutations in residues such as Asp116 or Glu119 on hGH, which contact GHR Chain C, can disrupt the receptor dimerization process. Even if the initial binding to the first receptor occurs, the failure to recruit and bind the second receptor molecule will prevent full receptor activation and downstream signaling. This can also lead to growth hormone resistance.

Mutations located in the loop regions or outside the main helical structures might affect protein folding, stability, or the subtle conformational changes necessary for optimal receptor binding. For instance, mutations near or within the disulfide bonds (C53-C165 and C182-C189) could disrupt the proper tertiary structure of GH, indirectly affecting its interaction with the GHR. The diversity and widespread distribution of the identified mutations underscore the complexity of genetic factors influencing growth. While some mutations may lead to a complete absence of functional GH, others might result in the secretion of a structurally altered hormone with impaired receptor binding or signaling capabilities.

The interaction of GH with its receptor involves a two-site binding mechanism, where one molecule of GH sequentially binds to two receptor monomers, leading to receptor dimerization and activation. Our contact analysis reveals the distinct sets of amino acid residues on GH that mediate these two binding events. The first interface (GH-GHR chain B) involves a broad range of interactions across different helical regions of hGH. The second interface (GH-GHR chain C) involves a separate set of residues, particularly in the N-terminal region and helix C of GH. The presence of several predicted functional (‘f’) and structural (‘s’) residues within these conserved regions further emphasizes their importance. Functional residues, being highly conserved and exposed, are likely involved in direct interactions, such as receptor binding. Structural residues, being highly conserved and buried, are crucial for maintaining the protein’s three-dimensional fold and stability, which indirectly supports proper receptor interaction.

The variability observed in some loop regions and the C-terminus might indicate that these regions are less critical for the core function of receptor binding or may be involved in more species-specific roles or interactions that do not necessitate high conservation across all orthologs. The presence of common mutations at the residues involved in both interfaces highlights the critical role of these regions in receptor binding and function. For example, mutations like R16C/L/H are located at a key residue that interacts with both receptor chains (Glu44 on chain C and is in proximity to the interface with chain B as seen in Figure 4). This suggests that mutations at this position are highly likely to disrupt the formation of the functional receptor dimer, further highlighting evolutionary pressure to maintain the integrity of these functional regions.

Similarly, mutations in the N-terminal region (e.g., P2Q, I4T/V, R8K, N12H, L15F) affect the second binding site, potentially impairing the initial engagement or the subsequent dimerization step. Mutations in helix C (e.g., G120C/S, D116N/E, E119D, T123M) also disrupt the second interface. The fact that different sets of mutations affect the two binding sites could have implications for the specificity and affinity of hGH for its receptor. Mutations affecting the first, higher-affinity binding site might have a more profound effect on overall receptor activation. However, disruptions in the second binding site, crucial for receptor dimerization, can also lead to significant impairments in growth hormone signaling. The strong overlap between the contact residues identified in this table and our contact analysis reinforces the accuracy of the interaction mapping.

The observation that many of the GH residues involved in contacting the receptor are highly conserved (as per ConSurf analysis) underscores the evolutionary pressure to maintain these crucial binding interfaces. These conserved residues likely play critical roles in the affinity and specificity of the GH-GHR interaction, which is essential for proper growth hormone signaling. The significant number of disease-associated mutations occurring at these contact residues highlights the direct link between disruptions in receptor binding and the development of growth disorders. Mutations in these key residues can potentially alter the conformation of GH at the binding interface, reduce its affinity for the receptor, or impair the formation of the functional receptor dimer.

Interestingly, several mutations predicted to significantly destabilize the GH protein by the SDM tool (Table x) also occur at these contact residues (e.g., P2Q, I4T, A17T, E119D, Y164H, K172N, E174K, I179V, C182R, R183H, C189Y, G190S). This suggests that the mechanism by which these mutations cause disease might involve both a disruption of the direct interaction with the receptor and a destabilization of the overall protein structure. Conversely, some mutations predicted to increase stability also occur at contact residues (e.g., N12H, H21Y, A24T, F25Y, S62C, Q91L, D112H, D116E, T123M, C189Y, G120C/S). While seemingly counterintuitive, increased stability could potentially hinder the conformational changes required for optimal receptor activation or release after binding.

The analysis of the ConSurf conservation scores in relation to the selected GH mutations reveals a trend where many disease-associated mutations occur at residues that are highly conserved across species. This observation supports the idea that these conserved residues play crucial roles in the structure, function, or interactions of the growth hormone protein. Mutations at such positions are more likely to disrupt these critical aspects, leading to phenotypic consequences such as growth hormone deficiency.

However, the presence of disease-associated mutations at less conserved positions, such as L15F and D112G/H, indicates that even variable residues can play a role in normal protein function, and changes at these sites can sometimes lead to disease. It is possible that these residues contribute to more subtle aspects of protein function or might be involved in interactions that are not universally conserved across all orthologs examined in the ConSurf analysis. The residue variety information provides additional context to the conservation scores. Positions with high conservation and low residue variety, like Cys53, likely have very specific structural or functional roles that cannot be easily substituted by other amino acids. Conversely, positions with low conservation and high residue variety, like Leu15, might be more tolerant to amino acid changes.

The results of the SDM analysis provide insights into the potential impact of genetic variations on the stability of the human growth hormone protein. Protein stability is crucial for proper folding, function, and interactions, and alterations in stability can lead to impaired biological activity and disease. The prediction that several disease-associated mutations destabilize the GH protein (e.g., L162P, A24T, L45P, A17T, E119D, Y164H) suggests that these mutations might disrupt the native conformation of the hormone, potentially affecting its binding affinity to the growth hormone receptor or its downstream signaling capabilities. For instance, L162P, predicted to cause the most significant destabilization, is located in helix D, a region known to be critical for receptor interaction (as seen in previous contact analysis). The introduction of a proline residue in a helical region can often disrupt the helix structure, leading to instability. Conversely, the prediction that some mutations increase protein stability (e.g., S79C, D116E, T123M, C182R, C189Y) is also interesting. While destabilizing mutations are often linked to loss-of-function phenotypes, increased stability could potentially affect the dynamics of the protein or its ability to undergo conformational changes necessary for receptor activation [1, 55].

It is important to note that changes in stability are just one aspect of how mutations can affect protein function. Other factors, such as changes in surface charge, hydrophobicity, or specific interactions with the receptor, can also play significant roles. For example, the R16C mutation, previously noted to reduce secretion, is predicted to be destabilizing by the SDM tool. Similarly, the R103C mutation in Kowarski syndrome, reported to have no effect on GHR signaling but stronger interaction with GHBP, shows a slightly destabilizing trend in this SDM analysis. Integrating the SDM analysis results with our previous findings on sequence conservation (ConSurf analysis) and known disease-causing mutations can provide a more comprehensive understanding of the molecular basis of growth hormone deficiencies. For example, mutations occurring in highly conserved residues that are also predicted to significantly destabilize the protein might be strong candidates for causing severe phenotypes [56].

Understanding the specific impact of each mutation on the two-site binding mechanism will contribute significantly to our knowledge of the molecular pathogenesis of growth disorders and may inform the development of targeted therapies. Further experimental studies by in vitro mutagenesis and functional assays, would be needed to validate the predictions and fully elucidate the impact of these mutations on growth hormone function and their association with growth disorders.

## 5. Conclusions

We have performed an analysis the mutations in human growth hormone by combining structural, evolutionary, and clinical data. The study highlights key amino acids that are essential for GH to bind to GHR, a crucial interaction for growth and metabolic processes. These amino acids are conserved across various vertebrate species, emphasizing their significance in GH function. We also found that disease-causing mutations frequently occur at these critical amino acid sites, disrupting the growth hormone’s ability to bind to its receptor and leading to growth disorders. The findings of this study offer a valuable framework for understanding the workings of human GH and the molecular underpinnings of growth hormone deficiency.

## Author Contributions

Conceptualization, A.V.P.; methodology, S.V. and A.V.P.; software, S.V. and A.V.P; validation S.V. and A.V.P.; formal analysis, S.V. and A.V.P.; investigation, AS.V.; resources, A.V.P..; data curation, S.V. and A.V.P.; writing—original draft preparation, S.V.; writing—review and editing, A.V.P.; visualization, S.V. and A.V.P.; supervision, A.V.P.; project administration, A.V.P.; funding acquisition, A.V.P. All authors have read and agreed to the published version of the manuscript.

## Institutional Review Board Statement

Not applicable.

## Informed Consent Statement

Not applicable.

## Data Availability Statement

All relevant data are provided in the manuscript.

## Abbreviations

The following abbreviations are used in this manuscript:

GH: Growth hormone
GHR: Growth hormone receptor
GHD: Growth hormone deficiency
GHBP: Growth hormone binding protein

## References

1. Rojas Velazquez, M. N.; Noebauer, M.; Pandey, A. V., Loss of Protein Stability and Function Caused by P228L Variation in NADPH-Cytochrome P450 Reductase Linked to Lower Testosterone Levels. International journal of molecular sciences 2022, 23, (17), 10141.

2. Prado, M. J.; Singh, S.; Ligabue-Braun, R.; Meneghetti, B. V.; Rispoli, T.; Kopacek, C.; Monteiro, K.; Zaha, A.; Rossetti, M. L. R.; Pandey, A. V., Characterization of Mutations Causing CYP21A2 Deficiency in Brazilian and Portuguese Populations. Int J Mol Sci 2021, 23, (1).

3. Parween, S.; Rojas Velazquez, M. N.; Udhane, S. S.; Kagawa, N.; Pandey, A. V., Variability in Loss of Multiple Enzyme Activities Due to the Human Genetic Variation P284T Located in the Flexible Hinge Region of NADPH Cytochrome P450 Oxidoreductase. Frontiers in Pharmacology 2019, 10, (1187).

4. Parween, S.; DiNardo, G.; Baj, F.; Zhang, C.; Gilardi, G.; Pandey, A. V., Differential effects of variations in human P450 oxidoreductase on the aromatase activity of CYP19A1 polymorphisms R264C and R264H. J Steroid Biochem Mol Biol 2020, 196, 105507.

5. Parween, S.; Fernandez-Cancio, M.; Benito-Sanz, S.; Camats, N.; Rojas Velazquez, M. N.; Lopez-Siguero, J. P.; Udhane, S. S.; Kagawa, N.; Fluck, C. E.; Audi, L.; Pandey, A. V., Molecular Basis of CYP19A1 Deficiency in a 46,XX Patient With R550W Mutation in POR: Expanding the PORD Phenotype. J Clin Endocrinol Metab 2020, 105, (4).

6. Alatzoglou, K. S.; Webb, E. A.; Le Tissier, P.; Dattani, M. T., Isolated growth hormone deficiency (GHD) in childhood and adolescence: recent advances. Endocr Rev 2014, 35, (3), 376–432.

7. Pandey, A. V., Bioinformatics tools and databases for the study of human growth hormone. Endocr Dev 2012, 23, 71–85.

8. Procter, A. M.; Phillips, J. A., 3rd; Cooper, D. N., The molecular genetics of growth hormone deficiency. Human genetics 1998, 103, (3), 255–72.

9. Hirt, H.; Kimelman, J.; Birnbaum, M. J.; Chen, E. Y.; Seeburg, P. H.; Eberhardt, N. L.; Barta, A., The human growth hormone gene locus: structure, evolution, and allelic variations. DNA 1987, 6, 59–70.

10. Mullis, P. E., Genetic control of growth. Eur J Endocrinol 2005, 152, (1), 11–31.

11. Phillips, J. A., 3rd; Cogan, J. D., Genetic basis of endocrine disease. 6. Molecular basis of familial human growth hormone deficiency. J Clin Endocrinol Metab 1994, 78, (1), 11–6.

12. de Vos, A. M.; Ultsch, M.; Kossiakoff, A. A., Human growth hormone and extracellular domain of its receptor: crystal structure of the complex. Science 1992, 255, (5042), 306–12.

13. Bluet-Pajot, M. T.; Epelbaum, J.; Gourdji, D.; Hammond, C.; Kordon, C., Hypothalamic and hypophyseal regulation of growth hormone secretion. Cellular and molecular neurobiology 1998, 18, (1), 101–23.

14. Kojima, M.; Hosoda, H.; Date, Y.; Nakazato, M.; Matsuo, H.; Kangawa, K., Ghrelin is a growth-hormone-releasing acylated peptide from stomach. Nature 1999, 402, (6762), 656–60.

15. Mullis, P. E.; Deladoey, J.; Dannies, P. S., Molecular and cellular basis of isolated dominant-negative growth hormone deficiency, IGHD type II: insights on the secretory pathway of peptide hormones. Horm Res 2002, 58, (2), 53–66.

16. Petkovic, V.; Miletta, M. C.; Eble, A.; Iliev, D. I.; Binder, G.; Fluck, C. E.; Mullis, P. E., Effect of zinc binding residues in growth hormone (GH) and altered intracellular zinc content on regulated GH secretion. Endocrinology 154, (11), 4215–25.

17. Norrelund, H., The metabolic role of growth hormone in humans with particular reference to fasting. Growth Horm IGF Res 2005, 15, (2), 95–122.

18. Meinhardt, U. J.; Ho, K. K., Modulation of growth hormone action by sex steroids. Clin Endocrinol (Oxf) 2006, 65, (4), 413–22.

19. Van Cauter, E.; Latta, F.; Nedeltcheva, A.; Spiegel, K.; Leproult, R.; Vandenbril, C.; Weiss, R.; Mockel, J.; Legros, J. J.; Copinschi, G., Reciprocal interactions between the GH axis and sleep. Growth Horm IGF Res 2004, 14 Suppl A, S10–7.

20. Widdowson, W. M.; Healy, M. L.; Sonksen, P. H.; Gibney, J., The physiology of growth hormone and sport. Growth Horm IGF Res 2009, 19, (4), 308–19.

21. Walenkamp, M. J.; Wit, J. M., Genetic disorders in the GH IGF-I axis in mouse and man. Eur J Endocrinol 2007, 157 Suppl 1, S15–26.

22. Clark, R. G.; Mortensen, D. L.; Carlsson, L. M.; Spencer, S. A.; McKay, P.; Mulkerrin, M.; Moore, J.; Cunningham, B. C., Recombinant human growth hormone (GH)-binding protein enhances the growth-promoting activity of human GH in the rat. Endocrinology 1996, 137, (10), 4308–15.

23. Leung, D. W.; Spencer, S. A.; Cachianes, G.; Hammonds, R. G.; Collins, C.; Henzel, W. J.; Barnard, R.; Waters, M. J.; Wood, W. I., Growth hormone receptor and serum binding protein: purification, cloning and expression. Nature 1987, 330, (6148), 537–43.

24. Niall, H. D., Revised primary structure for human growth hormone. Nat New Biol 1971, 230, (11), 90–1.

25. Hartman, M. L.; Faria, A. C.; Vance, M. L.; Johnson, M. L.; Thorner, M. O.; Veldhuis, J. D., Temporal structure of in vivo growth hormone secretory events in humans. The American journal of physiology 1991, 260, (1 Pt 1), E101–10.

26. Brown, R. J.; Adams, J. J.; Pelekanos, R. A.; Wan, Y.; McKinstry, W. J.; Palethorpe, K.; Seeber, R. M.; Monks, T. A.; Eidne, K. A.; Parker, M. W.; Waters, M. J., Model for growth hormone receptor activation based on subunit rotation within a receptor dimer. Nat Struct Mol Biol 2005, 12, (9), 814–21.

27. Gent, J.; Van Den Eijnden, M.; Van Kerkhof, P.; Strous, G. J., Dimerization and signal transduction of the growth hormone receptor. Mol Endocrinol 2003, 17, (5), 967–75.

28. Herrington, J.; Carter-Su, C., Signaling pathways activated by the growth hormone receptor. Trends Endocrinol Metab 2001, 12, (6), 252–7.

29. Chesover, A. D.; Dattani, M. T., Evaluation of growth hormone stimulation testing in children. Clin Endocrinol (Oxf) 2016, 84, (5), 708–14.

30. Cerbone, M.; Dattani, M. T., Progression from isolated growth hormone deficiency to combined pituitary hormone deficiency. Growth Horm IGF Res 2017, 37, 19–25.

31. Murray, P. G.; Dattani, M. T.; Clayton, P. E., Controversies in the diagnosis and management of growth hormone deficiency in childhood and adolescence. Arch Dis Child 2016, 101, (1), 96–100.

32. Petkovic, V.; Godi, M.; Pandey, A. V.; Lochmatter, D.; Buchanan, C. R.; Dattani, M. T.; Eblé, A.; Flück, C. E.; Mullis, P. E., Growth hormone (GH) deficiency type II: a novel GH-1 gene mutation (GH-R178H) affecting secretion and action. J Clin Endocrinol Metab 2010, 95, (2), 731–9.

33. Petkovic, V.; Eblé, A.; Pandey, A. V.; Betta, M.; Mella, P.; Flück, C. E.; Buzi, F.; Mullis, P. E., A novel GH-1 gene mutation (GH-P59L) causes partial GH deficiency type II combined with bioinactive GH syndrome. Growth Horm IGF Res 2011, 21, (3), 160–6.

34. Petkovic, V.; Miletta, M. C.; Boot, A. M.; Losekoot, M.; Flück, C. E.; Pandey, A. V.; Eblé, A.; Wit, J. M.; Mullis, P. E., Short stature in two siblings heterozygous for a novel bioinactive GH mutant (GH-P59S) suggesting that the mutant also affects secretion of the wild-type GH. Eur J Endocrinol 2013, 168, (3), K35–43.

35. Miletta, M. C.; Eblé, A.; Janner, M.; Parween, S.; Pandey, A. V.; Flück, C. E.; Mullis, P. E., IGHD II: A Novel GH-1 Gene Mutation (GH-L76P) Severely Affects GH Folding, Stability, and Secretion. J Clin Endocrinol Metab 2015, 100, (12), E1575–83.

36. Webb, E. A.; Dattani, M. T., Diagnosis of growth hormone deficiency. Endocr Dev 2010, 18, 55–66.

37. Franca, M. M.; Jorge, A. A.; Alatzoglou, K. S.; Carvalho, L. R.; Mendonca, B. B.; Audi, L.; Carrascosa, A.; Dattani, M. T.; Arnhold, I. J., Absence of GH-releasing hormone (GHRH) mutations in selected patients with isolated GH deficiency. J Clin Endocrinol Metab 2011, 96, (9), E1457–60.

38. Alatzoglou, K. S.; Kular, D.; Dattani, M. T., Autosomal Dominant Growth Hormone Deficiency (Type II). Pediatr Endocrinol Rev 2015, 12, (4), 347–55.

39. Altschul, S. F.; Madden, T. L.; Schaffer, A. A.; Zhang, J.; Zhang, Z.; Miller, W.; Lipman, D. J., Gapped BLAST and PSI-BLAST: a new generation of protein database search programs. Nucleic Acids Res 1997, 25, (17), 3389–402.

40. Ashkenazy, H.; Erez, E.; Martz, E.; Pupko, T.; Ben-Tal, N., ConSurf 2010: calculating evolutionary conservation in sequence and structure of proteins and nucleic acids. Nucleic Acids Res 2010, 38, (Web Server issue), W529-33.

41. Krieger, E.; Darden, T.; Nabuurs, S. B.; Finkelstein, A.; Vriend, G., Making optimal use of empirical energy functions: force-field parameterization in crystal space. Proteins 2004, 57, (4), 678–83.

42. Magrane, M., UniProt Knowledgebase: a hub of integrated protein data. Database (Oxford) 2011, 2011, bar009.

43. Worth, C. L.; Preissner, R.; Blundell, T. L., SDM--a server for predicting effects of mutations on protein stability and malfunction. Nucleic Acids Res 2011, 39, (Web Server issue), W215-22.

44. Pandurangan, A. P.; Ochoa-Montano, B.; Ascher, D. B.; Blundell, T. L., SDM: a server for predicting effects of mutations on protein stability. Nucleic Acids Res 2017, 45, (W1), W229-W235.

45. Topham, C. M.; Srinivasan, N.; Blundell, T. L., Prediction of the stability of protein mutants based on structural environment-dependent amino acid substitution and propensity tables. Protein Eng 1997, 10, (1), 7–21.

46. Reumers, J.; Schymkowitz, J.; Rousseau, F., Using structural bioinformatics to investigate the impact of non synonymous SNPs and disease mutations: scope and limitations. BMC Bioinformatics 2009, 10 Suppl 8, S9.

47. Karczewski, K. J.; Weisburd, B.; Thomas, B.; Solomonson, M.; Ruderfer, D. M.; Kavanagh, D.; Hamamsy, T.; Lek, M.; Samocha, K. E.; Cummings, B. B.; Birnbaum, D.; Daly, M. J.; MacArthur, D. G., The ExAC browser: displaying reference data information from over 60 000 exomes. Nucleic Acids Res 2017, 45, (D1), D840-D845.

48. Auton, A.; Brooks, L. D.; Durbin, R. M.; Garrison, E. P.; Kang, H. M.; Korbel, J. O.; Marchini, J. L.; McCarthy, S.; McVean, G. A.; Abecasis, G. R., A global reference for human genetic variation. Nature 2015, 526, (7571), 68–74.

49. Millar, D. S.; Lewis, M. D.; Horan, M.; Newsway, V.; Easter, T. E.; Gregory, J. W.; Fryklund, L.; Norin, M.; Crowne, E. C.; Davies, S. J.; Edwards, P.; Kirk, J.; Waldron, K.; Smith, P. J.; Phillips, J. A., 3rd; Scanlon, M. F.; Krawczak, M.; Cooper, D. N.; Procter, A. M., Novel mutations of the growth hormone 1 (GH1) gene disclosed by modulation of the clinical selection criteria for individuals with short stature. Human mutation 2003, 21, (4), 424–40.

50. Miyata, I.; Cogan, J. D.; Prince, M. A.; Kamijo, T.; Ogawa, M.; Phillips, J. A., Detection of Growth Hormone Gene Defects by Dideoxy Fingerprinting (ddF). Endocr J 1997, 44, (1), 149–154.

51. Deladoey, J.; Stocker, P.; Mullis, P. E., Autosomal dominant GH deficiency due to an Arg183His GH-1 gene mutation: clinical and molecular evidence of impaired regulated GH secretion. J Clin Endocrinol Metab 2001, 86, (8), 3941–7.

52. Takahashi, Y.; Kaji, H.; Okimura, Y.; Goji, K.; Abe, H.; Chihara, K., Brief report: short stature caused by a mutant growth hormone. N Engl J Med 1996, 334, (7), 432–6.

53. Petkovic, V.; Besson, A.; Thevis, M.; Lochmatter, D.; Eble, A.; Fluck, C. E.; Mullis, P. E., Evaluation of the biological activity of a growth hormone (GH) mutant (R77C) and its impact on GH responsiveness and stature. J Clin Endocrinol Metab 2007, 92, (8), 2893–901.

54. Takahashi, Y.; Shirono, H.; Arisaka, O.; Takahashi, K.; Yagi, T.; Koga, J.; Kaji, H.; Okimura, Y.; Abe, H.; Tanaka, T.; Chihara, K., Biologically inactive growth hormone caused by an amino acid substitution. J Clin Invest 1997, 100, (5), 1159–65.

55. Rojas Velazquez, M. N.; Therkelsen, S.; Pandey, A. V., Exploring Novel Variants of the Cytochrome P450 Reductase Gene (POR) from the Genome Aggregation Database by Integrating Bioinformatic Tools and Functional Assays. Biomolecules 2023, 13, (12), 1728.

56. Prado, M. J.; Ligabue-Braun, R.; Zaha, A.; Rossetti, M. L. R.; Pandey, A. V., Variant predictions in congenital adrenal hyperplasia caused by mutations in CYP21A2. Front Pharmacol 2022, 13, 931089.

